# Aberrant chromatin remodeling influences human neural cell fate change in Trisomy 21

**DOI:** 10.64898/2026.06.01.729170

**Authors:** Jenny A. Klein, Suraj Upadhya, Anna Nathanson, Dana Silvian, Eric F. Zaniewski, Robert Morris, Wilhelm Haas, Lindy E. Barrett

## Abstract

Correct neural progenitor cell (NPC) fate specification is essential to produce the full complement of neurons and glia needed for proper brain structure and function. Neurodevelopmental disorders, including the autosomal aneuploidy Down syndrome (DS), or Trisomy 21 (T21), are frequently associated with impaired cell fate decisions which ultimately drive differences in overall brain size and cell type composition through unknown mechanisms. To uncover mechanisms driving altered NPC fate in T21, we leverage paired single-nuclei transcriptomic and epigenomic analyses of human induced pluripotent stem cell (iPSC)-derived NPCs and their differentiated progeny coupled with in depth clonal cell fate, cell cycle, and proteomic analyses. Here we show that T21 NPCs fail to activate an orchestrated neurogenic program during the earliest stages of fate specification, instead maintaining a repressive chromatin structure over neurogenic loci, leading to reduced neurogenesis and continued NPC proliferation. We identify novel enrichment of the repressive histone mark H3K27me3 at fate instructive genes dysregulated across diverse cell and tissue types in T21, with corresponding genome-wide changes in H3K27me3 binding in T21 NPCs. Moreover, pharmacological treatment with an inhibitor of the Polycomb repressive complex 2 (PRC2) which catalyzes H3K27 methylation, is sufficient to partially restore neurogenesis in T21 cells. Collectively, our analyses reveal a chromatin mechanism influencing neurogenic defects in T21.

## INTRODUCTION

During mammalian corticogenesis *in vivo*, radial glial progenitor (RGP) cells undergo a period of expansion through symmetric cell divisions, after which they divide asymmetrically to first generate neurons followed by astrocyte generation^1,2^. This program is largely cell intrinsic, as RGPs maintain the same sequential differentiation potential *in vitro*^3^. Notably, rare mutations in genes such as *WDR62* or *ASPM,* as well as copy number variations (CNVs) including duplication at the 16p11.2 locus or the autosomal aneuploidies trisomy 21 (T21), trisomy 18 (T18), and trisomy 13 (T13), can drive brain undergrowth and differences in neural cell-type composition^4–7^. Of the CNVs, T21 or Down syndrome (DS) is currently the most well-characterized. Although underlying mechanisms remain unresolved, studies using human fetal brain tissue have identified a consistent reduction in neuron numbers from multiple brain regions in T21 at mid-gestation, with reduced brain size and cellularity which persist postnatally^8–10^. A recent single-cell analysis of fetal tissue from 13 to 23 weeks gestation showed a reduction in newborn neurons and increase in commitment to upper layer neuron identity at the expense of deep layer neuron identity, reflecting potential acceleration of the neurogenic timeline^11^. Other studies have also reported an increase in astrocyte number and fewer proliferating cells in T21^8–10^ all potentially contributing to altered neural composition. However, human neurogenesis begins within the first few gestational weeks, so the earliest stages of progenitor expansion and neurogenesis may not be fully captured in fetal tissue analyses. It remains to be determined the extent to which reduced neuron numbers observed in T21 are driven by increased cell death, decreased neurogenesis, and/or alterations in the temporal regulation of cortical development, such as precocious neurogenesis leading to progenitor pool depletion, or precocious astrogliogenesis.

Studies using human T21 induced pluripotent stem cell (iPSC) models consistently mirror reduced neuron output^12–15^ indicating that these models can be used to dissect underlying mechanisms and to capture the early neurogenic window that is difficult to acquire in tissue studies. Human iPSC models are particularly relevant to dissect this phenotype given the significant delay in neurogenesis observed in humans compared with other species, which facilitates cortical expansion^16^. However, studies in T21 have produced conflicting results when it comes to measurements of proliferation, cell death, and astrogliogenesis. For example, some studies report increased apoptotic cell death in T21^14,17^ while others report no change in cell death^15,18^. Studies focusing on comparative analyses across different dimensions (i.e., NPC subtype, time) support a complex picture. A comparison of interneuron progenitor subtypes found that overall proliferation rates remained unchanged in T21 for medial ganglionic eminence (MGE) progenitors but were reduced for caudal ganglionic eminence (CGE) progenitors^13^ suggestive of cell type differences. Looking over the time-course of NPC induction, another study reported reduced and increased proliferation rates at early and late stages, respectively^12^, reflecting the potential for dynamic cell cycle alterations over the course of cortical expansion. This complexity with regard to cell type and time-point is mirrored in fetal tissue analyses, where one study reports a reduction in apical progenitors in T21 over time, but increase in basal and intermediate progenitors at early to mid timepoints followed by a decrease in these populations at later time-points in T21^11^. Of note, T21 also drives dysregulation of cell fate in the hematopoietic system, where intrinsic differentiation biases, microenvironmental factors, additional genetic alterations, and proliferation changes have all been reported to contribute to cellular phenotypes^19–22^. The complexity of findings in T21 is perhaps not surprising when one considers aneuploidy from a cancer perspective; aneuploidy is a frequent event in cancer associated with poor prognosis, but also confers a proliferative disadvantage (the so-called “aneuploidy paradox”)^23,24^, underscoring the importance of cellular, genetic, and temporal context in the balance of growth and fate.

Here, we investigate the impacts of T21 on the earliest stages of human neural cell fate specification. Using independent isogenic T21/euploid iPSC pairs coupled with a forebrain NPC paradigm followed by spontaneous differentiation, we leverage clonal cell fate, cell cycle, proteomic, and single nuclei multiomic analyses to dissect underlying mechanisms. Our analyses reveal that at this early developmental stage, T21 drives reduced neurogenesis through profound re-wiring of repressive chromatin landscapes, with pharmacological inhibition of Polycomb repressive complex 2 (PRC2) sufficient to restore neurogenesis. These results identify for the first time, a chromatin pathway underlying early neural cell fate change in an autosomal aneuploidy.

## RESULTS

### Reduced neuron production from T21 NPCs

We generated NPCs from two independent isogenic T21/euploid iPSC pairs (XX: T21^WC^/Eupl^WC^ and XY: T21^DS^/Eupl^DS^)^25^ through the dual-SMAD inhibition strategy that expressed canonical NPC markers including PAX6 and SOX1 (**Fig. 1A and Fig. S1A-F**). As expected, transcriptomic analyses showed that NPCs most closely matched human prenatal time-points from the BrainSpan atlas with the highest correlation at 9-12 post conception weeks (**Fig. S1G**). We then allowed the NPCs to spontaneously differentiate using a minimal neural induction media (DMEM, N2, B27) to prevent extrinsic fate bias^26^. Spontaneous differentiation of NPCs unmasked a significant shift in cell fate between the T21 and isogenic euploid controls for both cell line pairs including in the number of neurons (MAP2+) and astrocyte precursor cells (CD44+, VIM+) generated (**Fig. S2**). The generation of astrocyte precursor cells is consistent with other studies using similar differentiation paradigms^17,26^. After one week of spontaneous differentiation, there were significantly fewer neurons produced by T21^WC^ NPCs compared to Eupl^WC^ NPCs (p=0.0015) (**Fig. 1B,C**). There was also a significant increase in the number of NPCs remaining in T21^WC^ compared to Eupl^WC^ (p=0.0064) and a trend toward increased astrocyte-precursor cells (p=0.89) (**Fig. 1B,C**). The T21^DS^/Eupl^DS^ isogenic pair showed the same cell fate phenotype with the T21^DS^ line producing significantly fewer neurons, more NPCs and a trend toward more astrocyte-precursors compared to the Eupl^DS^ line (p<0.0001;p<0.0001; p=0.99 respectively) (**Fig. 1D, E**). Thus, we observed fewer newborn neurons in T21.

**Figure 1.**
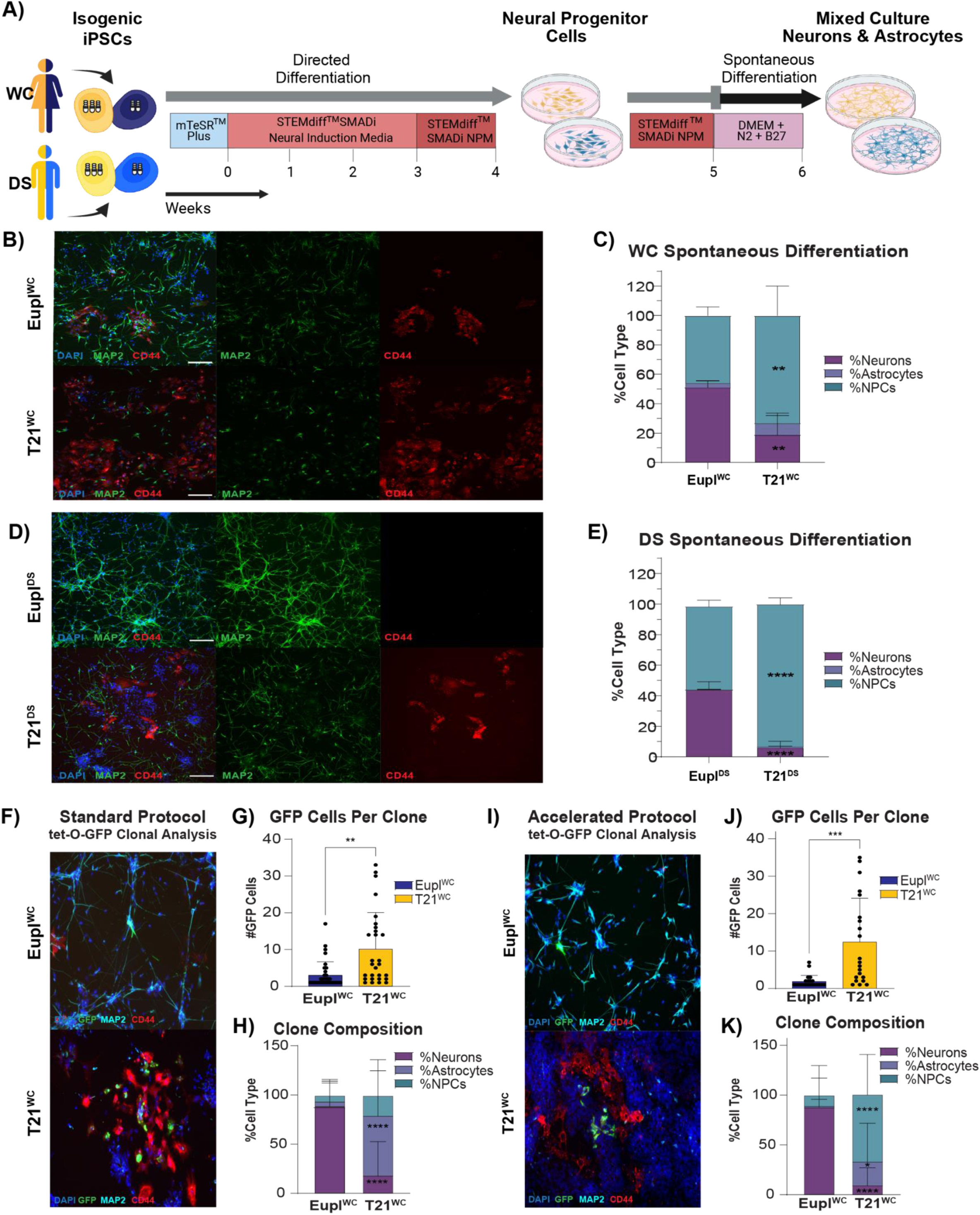
– Reduced neuron production from T21 NPCs. **A)** Schematic showing NPC generation followed by spontaneous differentiation to a mixed culture of neurons and astrocyte precursors (BioRender). **B)** Representative immunocytochemistry of T21^WC^/Eupl^WC^ cells after spontaneous differentiation showing neurons (MAP2+) and astrocyte precursors (CD44+), with DAPI counterstain. **C)** Significantly fewer neurons were produced by T21^WC^ (19.0 ± 14.6%) compared to Eupl^WC^ NPCs (51.3 ± 4.4%; p=0.0015). Significantly more NPCs remained in T21^WC^ compared to Eupl^WC^ (73.6 ± 20.0% vs 46.2 ± 5.8%; p=0.0064) and there was a trend toward increased astrocyte precursor cells in T21^WC^ (7.4 ± 5.5% in T21^WC^ vs 2.5 ± 1.7% in Eupl^WC^; p=0.89). **D)** Representative immunocytochemistry of the T21^DS^/Eupl^DS^ cells after spontaneous differentiation. **E)** T21^DS^ produced significantly fewer neurons (6.1 ± 4.2%) compared to Eupl^DS^ (44.1 ± 5.1%; p<0.0001). Significantly more T21^DS^ NPCs (93.5 ± 4.0%) remained compared to Eupl^DS^ (54.4 ± 4.0% NPCs, p<0.0001) and there was a trend towards more astrocyte precursors (0.4 ± 0.5% vs 0.06 ± 0.07% Eupl^DS^, p<0.0001). **F)** Representative immunocytochemistry of T21^WC^/Eupl^WC^ cells after spontaneous differentiation with sparse GFP labeling (1 in 10,000 GFP+ cells). **G)** Average number of cells produced by individual tetO-GFP Eupl^WC^ NPC clones (3.0 ± 3.7 cells) versus T21^WC^ NPC clones (10.1± 10.0 cells)(p=0.0011). **H)** Eupl^WC^ clones predominantly generated neurons (88.2 ± 26.0% neurons, vs 4.6 ± 19.4% astrocyte precursors and 7.2 ± 16.0% NPCs) while T21^WC^ clones produced significantly fewer neurons (17.4 ± 35.2% p<0.0001) and more astrocyte precursors and NPCs (61.0 ± 46.3% p<0.0001; 21.7 ± 35.8% p=0.19 respectively). **I)** Representative immunocytochemistry of T21^WC^/Eupl^WC^ cells after the accelerated spontaneous differentiation protocol (−1 week) with sparse GFP labeling. **J)** Average number of cells produced by individual tet-O-GFP Eupl^WC^ clones (1.9 ± 1.6 cells) versus tetO-GFP T21^WC^ clones (12.4 ± 11.7 cells) (p=0.0005). **K)** tet-O-GFP Eupl^WC^ clones predominantly generated neurons (87.2 ± 30.0%) compared to 1.5 ± 7.1 % astrocyte precursors and 11.3 ± 29.7% NPCs. tet-O-GFPT21^WC^ clones produced significantly fewer neurons compared with Eupl^WC^ (8.9 ± 18.1% p<0.0001) and significantly more astrocyte precursors (23.8 ± 39.1% p=0.046) and NPCs (67.3 ± 40.9% p<0.0001). Significance was determined by Two-way ANOVA followed by Sidak test for multiple comparisons for C), E), H), and K) and Welch’s t test for G) and J). A minimum of n=3 independent wells per condition per karyotype were used for each experiment. Each well was independently passaged and differentiated starting from the iPSC stage to generate biological replicates.

### Clonal analysis shows continued T21 NPC proliferation

The reduction in neuron number from T21 relative to the isogenic euploid conditions is consistent with previous studies from human postmortem tissue^8–10^ and iPSC-based models^12–15^, and could be due to several factors, including reduced neurogenesis from a comparable number of NPCs, a delay in the timing of neurogenesis (i.e., continued proliferation at the expense of differentiating divisions), premature neurogenesis leading to progenitor cell depletion, increased neuronal cell death or increased senescence. The increased number of undifferentiated NPCs in the T21 conditions relative to the euploid conditions (**Fig. 1B-E**) would argue against premature neurogenesis leading to progenitor cell depletion at this stage, based on the bulk assay. To further distinguish between these possibilities, we first used sparsely labeled tetO-GFP Eupl^WC^ and tetO-GFP T21^WC^ NPCs to perform clonal analyses (1:10,000 GFP+ to GFP-cells; **Fig. 1F-H, S3A**). Individual tetO-GFP Eupl^WC^ NPC clones produced significantly fewer cells during spontaneous differentiation than T21^WC^ NPC clones consistent with increased proliferation in the latter karyotype (p=0.0011) (**Fig. 1F-G**). As with bulk analyses from both isogenic pairs, T21^WC^ clones produced significantly fewer neurons (p<0.0001) and more astrocyte precursors and NPCs (p<0.0001 and p=0.19 respectively) compared with Eupl^WC^ clones, which predominantly generated neurons (**Fig. 1H**).

To assess precocious neurogenesis, we forced the NPCs to spontaneously differentiate one week early, relative to the standard protocol (**Fig. 1I-K, S3B**). Here, individual tetO-GFP Eupl^WC^ clones again produced significantly fewer cells than individual tetO-GFP T21^WC^ clones (p=0.0005), consistent with increased proliferation in the T21 condition (**Fig. 1J**). Eupl^WC^ clones still predominantly generated neurons while individual T21^WC^ clones produced significantly fewer neurons compared with Eupl^WC^ (p<0.0001) and significantly more astrocyte precursors (p=0.046) and NPCs (p<0.0001) (**Fig. 1K**). Thus, tetO-GFP T21^WC^ clones did not show increased neuron production with a shortened differentiation time-course, arguing against the propensity for precocious neurogenesis at this early time-point.

Moreover, in both the standard and accelerated differentiation paradigms, there was a significant correlation between the number of progeny produced and cell fate (**Fig. S3C-D**). In the standard time course, there was a negative correlation between the number of GFP+ cells and the percentage of neurons per clone (R^2^=0.2608, p<0.0001) (**Fig. S3C**). Conversely, the number of GFP+ cells was positively correlated with the percentage of astrocyte precursor cells per clone (R^2^=0.1151, p=0.0070) as well as the percentage of NPCs per clone (R^2^=0.1030, p<0.0001) (**Fig. S3C**). These correlations were also present in the accelerated differentiation paradigm, where there was again a negative correlation between the number of GFP+ cells and percentage of neurons per clone (R^2^=0.2667, p<0.0001) and a positive correlation between the number of GFP+ cells and the percentage of NPCs per clone (R^2^=0.3673, p<0.0001) (**Fig. S3D**).

Additionally, to rule out premature senescence or increased cell death as drivers of the reduced neuron numbers in T21, we performed immunohistochemical staining for cyclin dependent kinase inhibitor 2A (p16INK4A), a marker of senescence, and cleaved caspase 3 (CC3), a marker of apoptosis. Analysis of P3 NPCs showed no difference in either marker between conditions (**Fig. S3E-H**). Together, our data are consistent with continued NPC proliferation at the expense of neuron generation at the earliest stages of differentiation in T21.

### Alterations in the G1 / S ratio support continued proliferation of T21 NPCs

While NPC proliferation has been extensively studied in the context of T21 with variable findings reflecting the cell type, time point, and methodology^12–15,17,18^, to date, no explicit mechanistic link between altered cell proliferation and cell fate has been drawn in T21. To rigorously quantify cell cycle phenotypes in our human iPSC derived NPCs, we performed time-course analyses in multiple independent genetic backgrounds and used a series of orthogonal approaches to capture both temporal changes and multiple aspects of cell cycle, including shifts in proportion of cell cycle phase and length (**Fig. 2A-G, Fig. S4A-D, Fig. S5A-G**).

**Figure 2.**
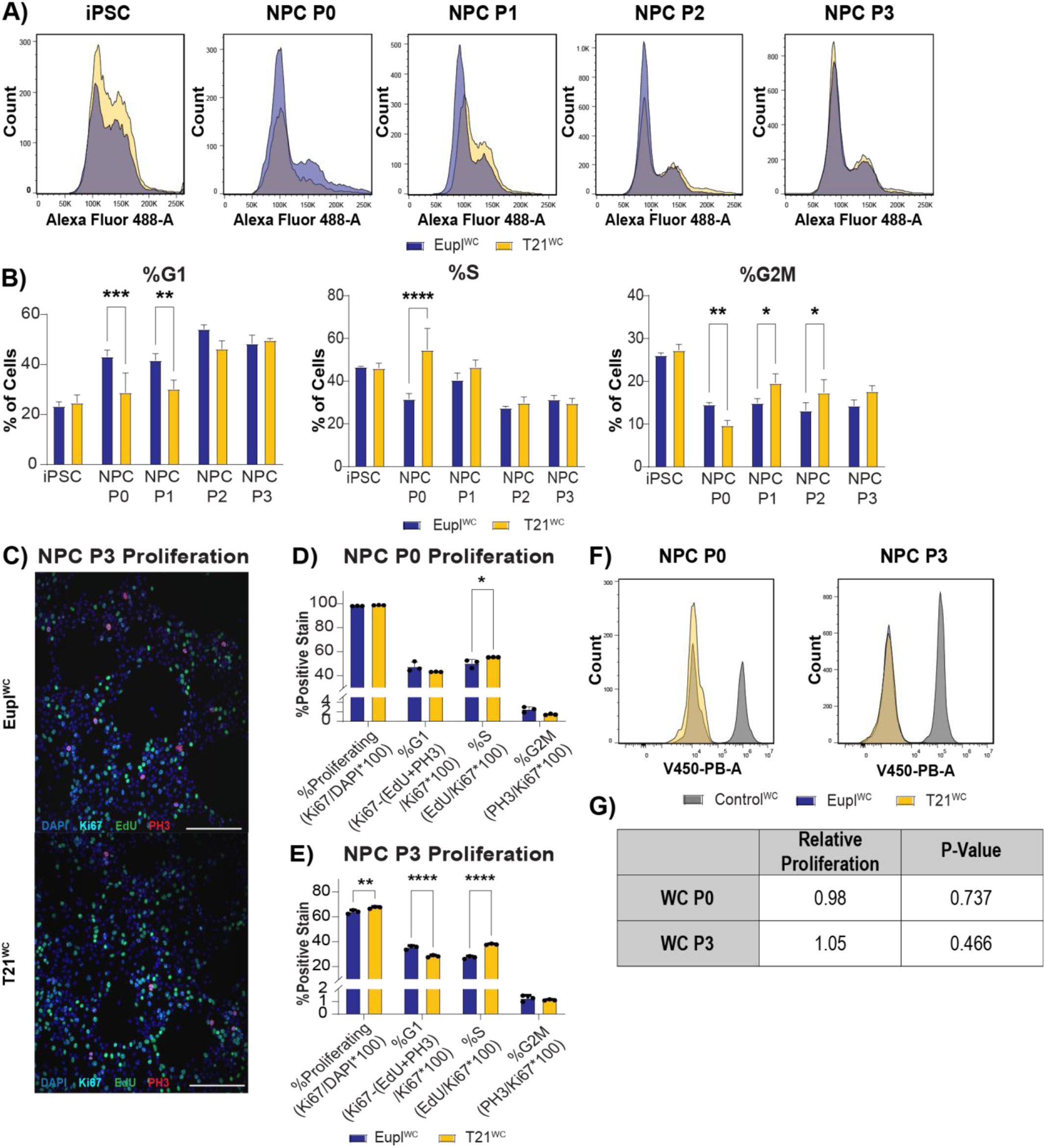
– Alterations in the G1 / S ratio support continued proliferation of T21 NPCs. **A)** Representative traces of flow cytometry analysis of Vybrant DyeCycle Green intensity over a five-week differentiation time course of Eupl^WC^ and T21^WC^ cells. **B)** Quantification of dye intensity identified significant shifts in all cell cycle phases between karyotypes. At P0, there was a significant decrease in the proportion of NPCs in G1 in T21^WC^ cell lines (43.1 ± 2.6% Eupl^WC^ vs 28.8 ± 7.8% in T21^WC^ p=0.022) with a concurrent increase in the proportion of NPCs in S phase (31.6 ± 2.6% Eupl^WC^ vs 54.7 ± 10.0% in T21^WC^ p=0.0007). This decrease in G1 relative to Eupl^WC^ NPCs continued to P1 (41.6 ± 2.7% Eupl^WC^ vs 30.2 ± 3.6% in T21^WC^ p=0.001). Modest but significant changes in the proportion of cells in G2M were identified (P0, p=0.0076), P1, p=0.0109, P2, p=0.0244). **C)** Representative immunocytochemistry of P3 NPCs immunostained for Ki67 (all cycling cells), EdU (cells in S phase), and PH3 (cells in G2/M phase). **D-E)** Quantification of cell cycle phases for P0 (D) and P3 (E) NPCs. At P0, there was a significant increase in the proportion of NPCs in S phase in T21^WC^ compared to Eupl^WC^ (50.1 ± 3.4% Eupl^WC^ vs 55.4 ± 0.1% in T21^WC^ p=0.015). At P3, there was a significant decrease in the proportion of T21 NPCs in G1 phase (G1: 35.4 ± 1.6% Eupl^WC^ vs 28.4 ± 0.9% in T21^WC^ p<0.0001) and a significant increase in the proportion in S phase (27.4 ± 1.1% Eupl^WC^ vs 38.0 ± 0.5% in T21^WC^ p<0.0001). At P3, there were significantly more proliferative NPCs overall in T21^WC^ (64.0 ± 1.5% Eupl^WC^ vs 67.5 ± 0.8% in T21^WC^ p=0.0021). **F)** Representative traces of flow cytometry analysis of CellTrace Violet in Eupl^WC^ and T21^WC^ NPCs. **G)** Quantification of CellTrace Violent results shown in (F). Significance was determined by Two-way ANOVA followed by Sidak test for multiple comparisons for B), D) and E) and by Welch’s t test for G). A minimum of n=3 independent wells per condition per karyotype were used for each experiment. Each well was independently passaged and differentiated starting from the iPSC stage to generate biological replicates.

We first assessed shifts in the proportion of cell cycle phases driven by T21 using flow cytometry analysis of Vybrant DyeCycle Green intensity over a five-week differentiation time course including iPSCs and P0 to P3 NPCs in both T21^WC^/Eupl^WC^ and T21^DS^/Eupl^DS^ (**Fig. 2A-B, Fig. S4A, Fig. S5A**). While T21 did not impact the proportion of cell cycle phases at the iPSC stage in either genetic background (**Fig. 2B, Fig. S5A**), significant changes emerged during NPC differentiation. At P0, there was a significant decrease in the proportion of NPCs in G1 in both T21 cell lines (WC p=0.022; DS p<0.0001) with a concurrent increase in the proportion of NPCs in S phase (WC p=0.0007; DS p<0.0001) (**Fig. 2B, Fig. S5A**). In the T21^WC^ NPCs, this decrease in G1 relative to Eupl^WC^ NPCs continued at P1 (p=0.001) (**Fig. 2B**). Additionally, modest but significant changes in the proportion of cells in G2M were also identified in both isogenic pairs (**Fig. 2B, Fig. S5A**).

While flow cytometry analyses identified shifts in G1/S proportions in T21, they do not account for differences in the total number of proliferating cells, which could influence the above proportions. We therefore used immunohistochemical staining to first identify all proliferating cells (Ki67) followed by combinatorial staining of EdU (S phase) and PH3 (G2M) (**Fig. 2C-E, Fig. S4C-D, Fig. S5B-E**). Using this methodology, we again identified a significant increase in the proportion of cells in S phase in P0 NPCs in both T21^WC^ and T21^DS^ compared to their isogenic euploid controls (WC p=0.015; DS p<0.0001) along with a decrease in the proportion of NPCs in G1 in T21^DS^ (p<0.0001) (**Fig. 2D, Fig. S5C**). Additionally, we identified this same shift in the G1/S ratio in P3 NPCs (G1 p<0.0001; S p<0.0001) (**Fig. 2E**) further confirming shifts in the G1/S ratio during NPC development in T21.

Multiple studies from *in vivo* models suggest a relationship between G1/S length and neurogenesis. G1 lengthening in mouse models has been associated with the transition to fate restricted basal intermediate progenitors, with expanding progenitors investing more time in S-phase than their committed neurogenic counterparts^27^. Reducing G1 length in mouse models has also been shown to delay neurogenesis and promote expansion of progenitor populations^28^, consistent with studies in *Drosophila* where G1 lengthening has been shown to promote neurogenesis^29^. Thus, the decreased G1 and increased S phase observed in T21 NPCs could reflect continued proliferation and reduced neurogenesis. Alternatively, decreased G1 and increased S phase has also been associated with aneuploidy driven replicative stress in other cellular systems^30,31^ which could lead to reduced cycling and fewer neurons. We therefore used CellTrace Violet (CTV) to assess cell cycle length and performed whole proteome analyses to globally assess cell cycle regulators. Analyses with CTV showed that at both P0 and P3, T21^WC^ and Eupl^WC^ NPCs proliferated at the same rate, indicating that the shift was in the ratio of G1 to S, rather than cells arresting in S in the T21 condition (**Fig. 2F,G**). Unexpectedly, T21^DS^ showed a significant decrease in cell cycle length compared with its isogenic euploid control (**Fig. S5F-G**), indicating faster proliferation, atypical for T21 NPCs. Consistent with this result, mass spectrometry based whole-proteome analyses of P3 NPCs from both cell line pairs **(Fig. S6A-B, Table S1)** revealed that T21^DS^ had significantly upregulated proteins associated with proliferation such as minichromosome maintenance 2, 5, and 7 (MCM2, MCM5, and MCM7) compared with Eupl^DS^, and in contrast to T21^WC^ (**Fig. S6C**). Combined, our results suggest that the T21^DS^ cell line may have made adaptations to aneuploidy not evident in the T21^WC^ cell line. Despite these adaptations, both T21 cell lines shared similar changes in G1/S ratios and cell fate phenotypes independent of differences in overall proliferation rates, and consistent with reduced neurogenesis.

### Multiomics reveals a loss of neurogenic gene expression programs in T21 NPCs

To identify molecular mechanisms driving altered neurogenesis in T21, we performed single nuclei 10X Multiome ATAC + Gene Expression sequencing on NPCs and spontaneously differentiated cells using the Eupl^WC^/T21^WC^ isogenic pair (**Fig. 3A-D**, **Fig. 4A-H, Fig. S7A-F, Fig. S8A-D**). Post sequencing, a total of 20,145 cells passed quality control metrics with 1,089 to 2,661 median genes detected per cell (**Fig. S7A**). RNA based clustering identified twelve distinct cell clusters ranging from cycling NPCs to different subtypes of excitatory and inhibitory neurons and astrocyte precursors (**Fig. 3A, Fig. S7E-F**). Similar to initial characterization (**Fig. 1C**), cellular composition varied by karyotype, with Eupl^WC^ generating more neurons relative to T21^WC^, and T21^WC^ generating more NPCs and astrocyte precursors relative to Eupl^WC^ (**Fig. 3B**).

**Figure 3.**
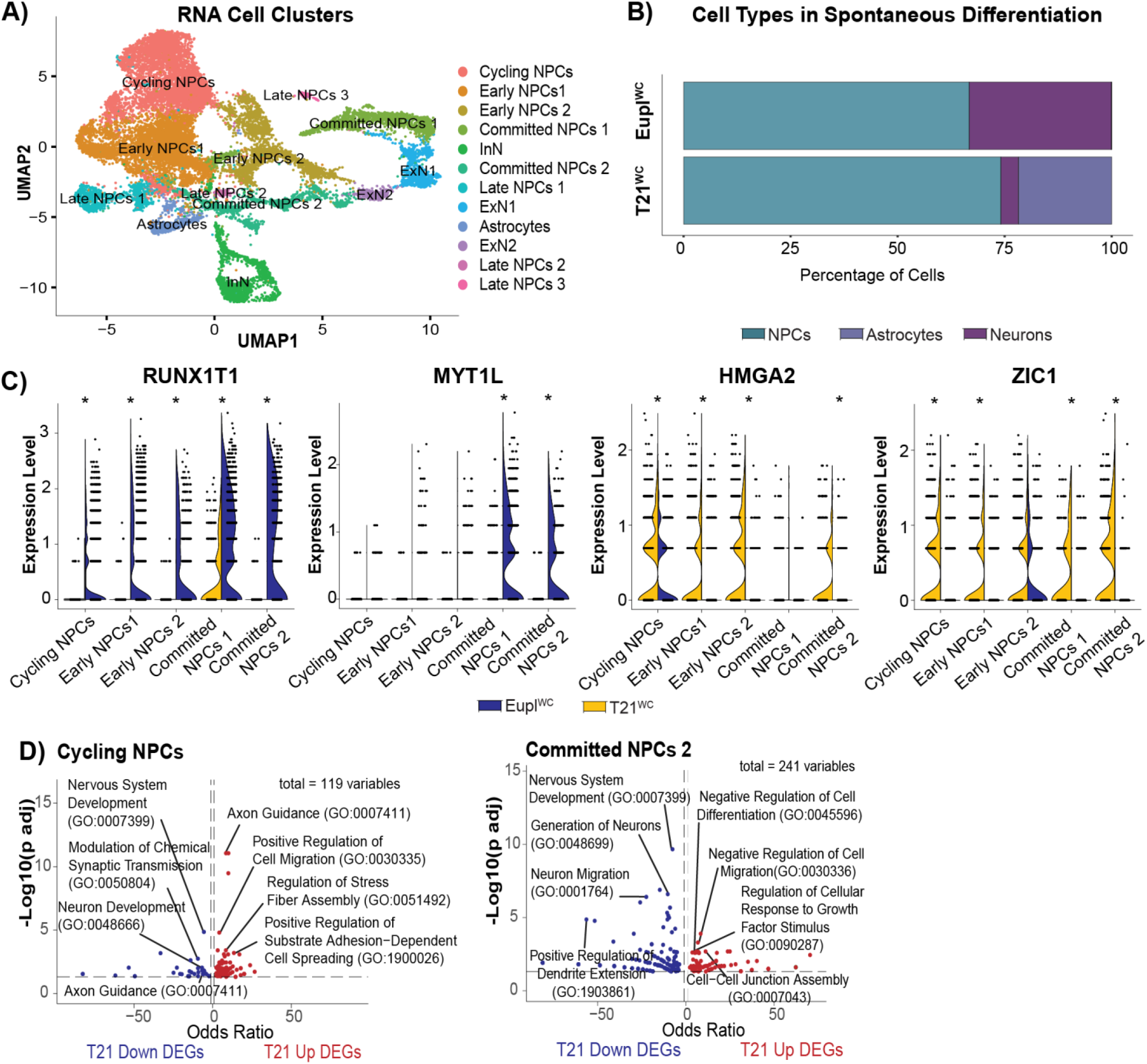
– Multiomics reveals a loss of neurogenic gene expression programs in T21 NPCs. **A)** UMAP clustering of Eupl^WC^ and T21^WC^ single nuclei based on their RNA profile produced 12 distinct clusters ranging from cycling NPCs to neurons and astrocyte precursors. **B)** Cellular composition for Eupl^WC^ and T21^WC^ cells. **C)** Examples of individual DEGs from snRNA-seq data pseudobulked per cluster including *RUNX1T1*, *MYT1L*, *HMGA2*, and *ZIC1*. Significance was determined by the two-sided Wilcoxon Rank-Sums test followed by Benjamini and Hochberg correction for multiple comparisons. Adjusted p-values of < 0.05 were considered significant. **D)** Gene ontology analysis from Cycling NPCs and Committed NPCs 2 clusters. A minimum of n=3 independent wells per condition per karyotype were used for each experiment. Each well was independently passaged and differentiated starting from the iPSC stage to generate biological replicates.

**Figure 4.**
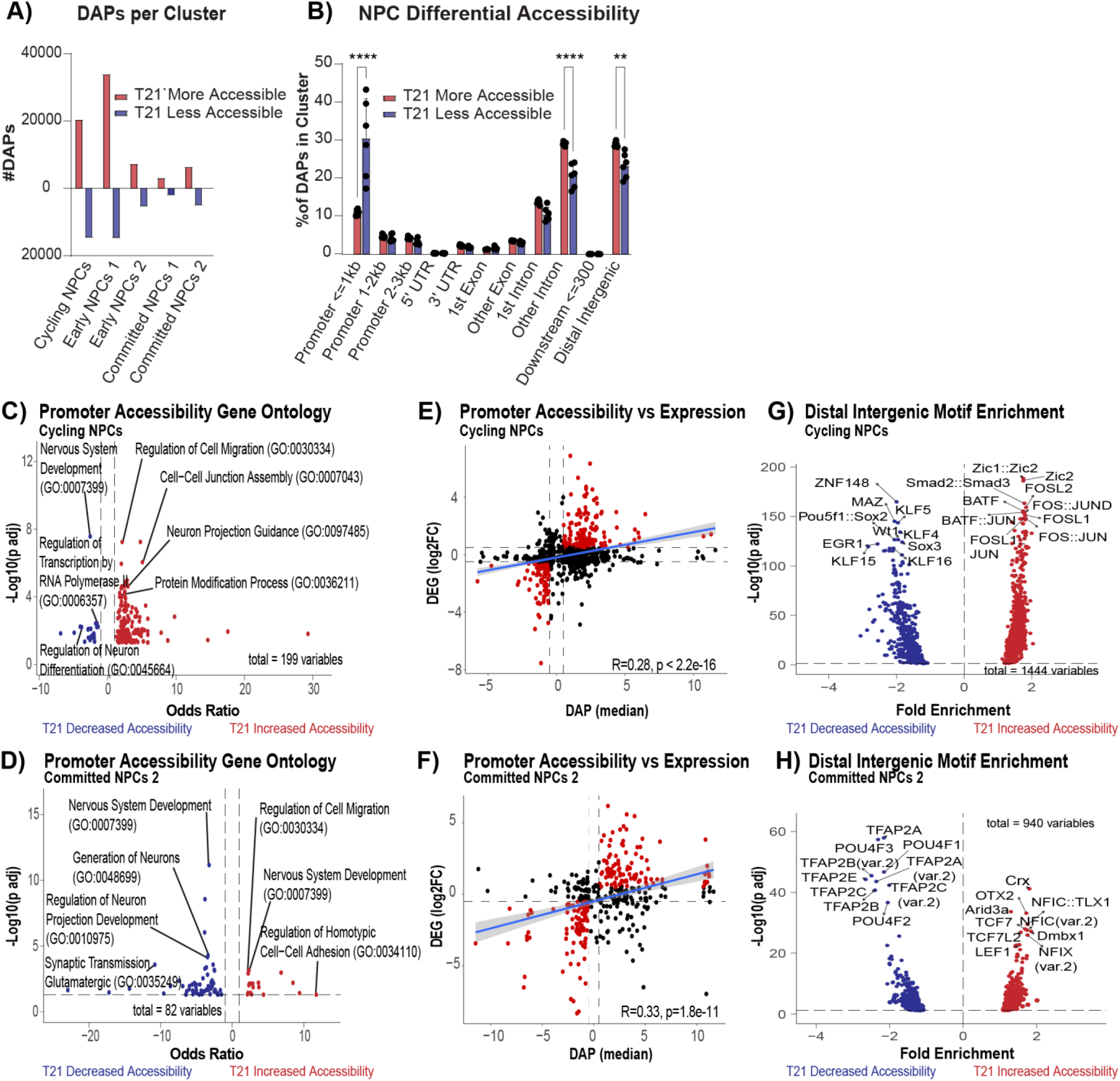
– T21 drives decreased chromatin accessibility at neurogenic loci. **A)** UMAP clustering of Eupl^WC^ and T21^WC^ single nuclei based on their ATAC profile overlaid with cell identity derived from RNA clusters. **A)** Number of differentially accessible peaks (DAPs) per cluster which gained or lost accessibility in T21^WC^ NPCs compared to Eupl^WC^ NPCs. **B)** Categorizing the DAPs by genomic location identified more decreased accessibility in T21^WC^ proximal promoters (30.7 ± 10.4% decreased accessibility vs 11.1 ± 0.8% increased accessibility, p<0.0001) and more increased accessibility in T21^WC^ non-coding regions compared to decreased accessibility (29.3 ± 0.5% increased intron accessibility vs 20.8 ± 3.1% decreased intron accessibility p<0.0001 and 29.1 ± 0.7% increased distal intergenic accessibility vs 23.5 ± 3.3% decreased distal intergenic accessibility p=0.0018). Significance was determined by Two-way ANOVA followed by Sidak test for multiple comparisons. **C-D)** Gene ontology analysis from DAPs in proximal promoters for Cycling NPCs (C) and Committed NPCs 2 (D). **E-F)** General linear regression between differential expression and differential promoter accessibility for Cycling NPCs (R=0.28; p<2.2e-16) (E) and Committed NPCs 2 (R=0.33; p=1.8e-11)(F). **G-H)** Motif enrichment analysis of distal intergenic regions for Cycling NPCs (G) and Committed NPCs 2 (H).

To identify molecular changes leading to altered cell fate in T21, we focused our analysis on the NPC clusters. Pseudobulked differential analysis identified hundreds of differentially expressed genes (DEGs) per cluster consistent with broad *trans* effects on gene expression (**Fig. 3C, Fig. S9A**). Indeed, from pseudobulked NPCs only 12 of the upregulated genes were encoded on HSA21 (**Fig. S10A**) and from whole proteome analyses only 22 of the upregulated proteins were encoded on HSA21 (**Fig. S10B**). Multiple genes known to play a role in neuronal fate, such as *RUNX1 partner transcriptional co-repressor 1* (*RUNX1T1)*, *Myelin transcription factor 1 like* (*MYT1L)*, *High mobility group AT-hook 2 (HMGA2)*, and *Zic family member 1 (ZIC1)* were found to be dysregulated in T21 (**Fig. 3C, Fig. S9A, Table S2**). *MYT1L* for example, is a pro-neuronal transcription factor important for suppressing non-neuronal cell fates^32^ which was upregulated during NPC differentiation in the Eupl^WC^ condition, but which failed to turn on in the T21^WC^ condition (**Fig. 3C, Table S2**). By contrast, *HMGA2* is a pro-proliferative chromatin factor^33^ which was expressed under both conditions in Cycling NPCs, but which failed to turn off in the T21^WC^ condition (**Fig. 3C, Table S2**). Gene ontology analysis of DEGs from each NPC cluster consistently identified neurogenesis terms such as Neuron Development, Neuron Differentiation, and Generation of Neurons as significantly overrepresented in the downregulated genes in T21 (**Fig. 3D, Fig. S9B**). By contrast, terms such as Negative Regulation of Cell Differentiation, Regulation of Cell Migration, and Cytoplasmic Translation were significantly overrepresented among the upregulated genes in T21 (**Fig. 3D, Fig. S9B**). Consistent with our transcript level findings, whole-proteome analyses of P3 NPCs also revealed decreased expression of neurogenic proteins in both the T21^WC^ and T21^DS^ conditions, relative to their isogenic euploid controls, although individual protein changes varied between cell lines (**Fig. S11**). Collectively, these data indicate dysregulation in the molecular programs driving NPCs towards a neural fate but do not implicate a single candidate factor.

### T21 drives decreased chromatin accessibility at neurogenic loci

As 10x Multiome sequencing produces complementary snATAC-seq profiles in parallel with snRNA-seq, we analyzed changes in chromatin accessibility in T21^WC^ and Eupl^WC^ NPCs. T21^WC^ NPCs showed broad changes in accessibility with both gains and losses of differentially accessible peaks (DAPs) across all NPC clusters (**Fig. 4A, Table S3**). Categorizing the DAPs by genomic location identified most of the changes in proximal promoters and noncoding regions, with more decreased accessibility in T21^WC^ proximal promoters compared to increased accessibility (p<0.0001) and more increased accessibility in T21^WC^ non-coding regions compared to decreased accessibility (intron accessibility p<0.0001; distal intergenic accessibility p=0.0018) (**Fig. 4B, Table S3**). The loss and gain of accessibility in other genomic regions was modest and balanced between gains and losses (**Fig. 4B, Table S3**).

Focusing on differential accessibility first at proximal promoters, gene ontology analysis from each NPC cluster consistently identified neurogenesis terms such as Regulation of Neuron Differentiation, Neuron Differentiation, and Nervous System Development as significantly overrepresented in the genes with decreased proximal promoter accessibility in T21 (**Fig. 4C-D, Fig. S9C**), paralleling the transcriptional data (**Fig. 3D, Fig. S9B**). By contrast, terms such as Regulation of Cell Migration, Extracellular Matrix Organization, and Protein Modification Process were significantly overrepresented among the genes with increased proximal promoter accessibility in T21 (**Fig. 4C-D, Fig. S9C**). Performing a linear regression on genes that showed both differential expression and differential promoter accessibility between Eupl^WC^ and T21^WC^ revealed modest but significant correlations between expression and accessibility in each of the NPC clusters (**Fig. 4E-F, Fig. S9D**). Analysis of distal intergenic regions revealed that restructuring of the regulatory landscape may also contribute to the neurogenic changes in T21. For example, cycling NPCs gained accessibility in regions enriched for Activator Protein-1 (AP-1) motifs previously associated with T21^34^, as well as other pro-proliferative factors like *Zic family member 2 (ZIC2),* while motifs associated with NPC fate specification and neural development like *Early growth response 1 (EGR1)* lost accessibility (**Fig. 4G**). Committed NPCs also lost accessibility in regions enriched for transcription factor AP-2 family member motifs and other neurogenic factors like the BRN-3 / POU family while gaining accessibility in regions enriched for Nuclear Factor I (NFI) motifs associated with astrocyte development (**Fig. 4H, Fig. S9E**). These data implicate a chromatin mechanism in the neurogenic gene expression changes in T21.

### T21 alters NPC and neuron cell density along developmental pseudotime

To understand how differences in expression and accessibility affect developmental trajectory we performed a pseudotime analysis to identify distinct developmental lineages. Setting cycling NPCs as the starting cluster, we identified seven lineages including three lineages that terminated in neuronal clusters, one that terminated in an astrocyte precursor cluster, and three that terminated in progenitor clusters (**Fig. 5A**).

**Figure 5.**
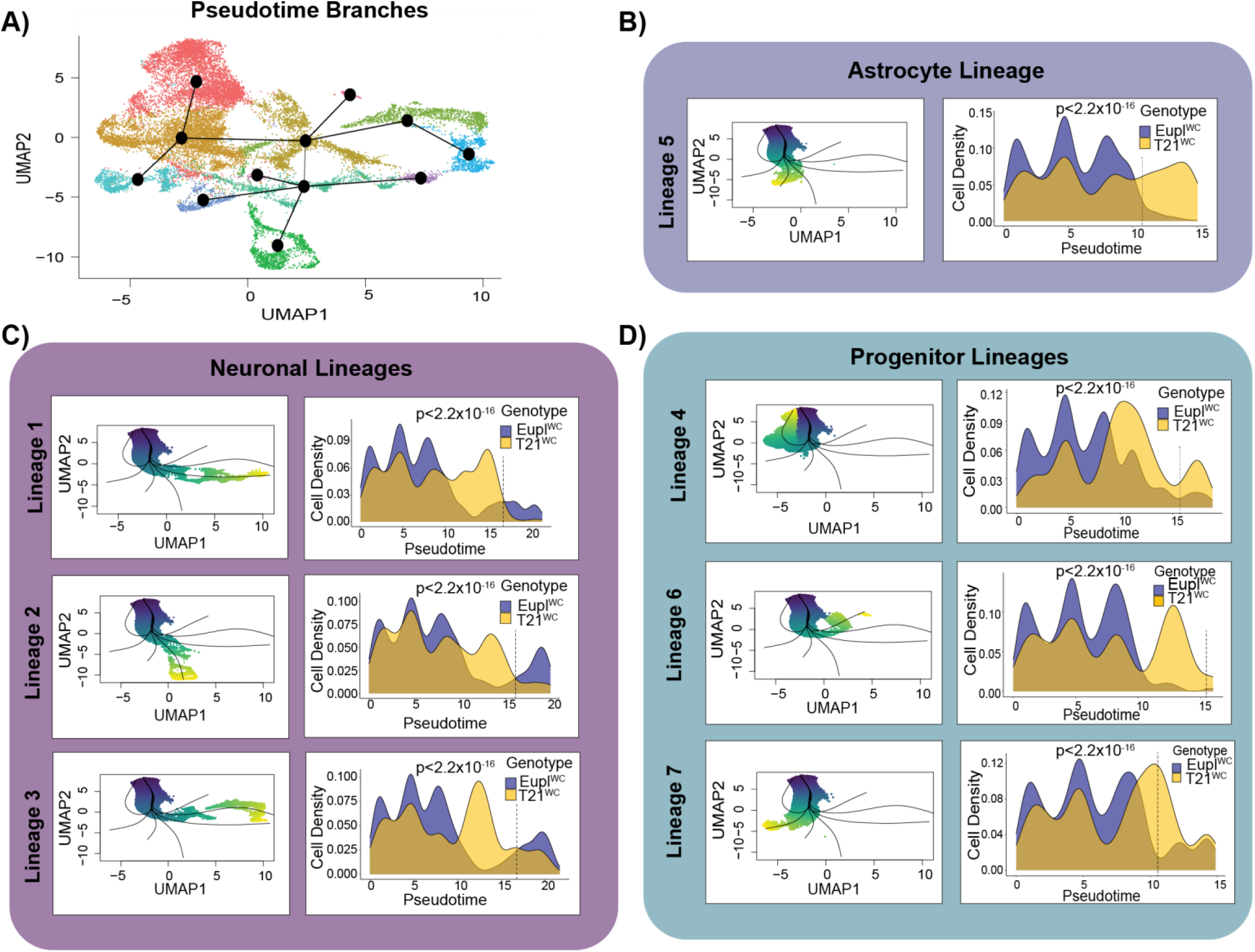
– T21 alters NPC and neuron cell density along developmental pseudotime. A) Pseudotime analysis identified seven developmental lineages of cells with the cycling NPC cluster set at the starting cluster. B-D) Cell density versus pseudotime values plotted between karyotypes for one astrocyte precursor lineage (B) three neuronal lineages (C) and three progenitor lineages. (D) Significance was determined by quasibinomial logistic regression and p<2.2×10^−16^ for each lineage. Dotted lines represent terminal differentiation.

To assess lineage progression, we first performed Kolmogorov-Smirnov tests on each lineage to identify statistical differences in mean pseudotime values between the Eupl^WC^ and T21^WC^ conditions. We found statistically significant differences in each of the seven lineages, with the T21^WC^ condition consistently showing a higher mean pseudotime value (**Fig. S12**), which could be interpreted as accelerated maturation compared to euploid. However, this analysis does not account for differences in distribution along the lineage. To assess differences in lineage trajectory with increased resolution, we separated each lineage by karyotype, plotted cell density versus pseudotime value, and subsequently performed general linear regression to identify significant differences in density distributions between karyotypes along each lineage (**Fig. 5 B-D**). Using this approach, Eupl^WC^ cells clearly had an increased density at higher pseudotime values in all three neuronal lineages compared with T21^WC^ (p<2.2e-16; **Fig. 5C**) while T21^WC^ cells showed an increased density at higher pseudotime values in the astrocyte precursors (p<2.2e-16; **Fig. 5B**) and NPC lineages (p<2.2e-16; **Fig. 5D**). Thus, increased density of T21^WC^ NPCs prior to differentiation (dashed line) in the neuronal lineages likely accounts for the higher cumulative mean pseudotime values in T21 as opposed to increased maturation. This result is also consistent with our clonal analyses using an accelerated differentiation paradigm, where the T21 NPCs still produced fewer neurons compared to isogenic euploid control neurons (**Fig. 1 I-K; Fig. S3A-D**), arguing against precocious neurogenesis in T21 at the earliest stages of differentiation.

### T21 dysregulates gene regulatory networks driving neurogenesis

Individually, our snRNA-seq and snATAC-seq data show decreased expression and promoter accessibility of neurogenic genes in T21 NPCs, consistent with the decreased neurogenesis. However, to take advantage of the power available from paired single nuclei RNA-seq and ATAC-seq data and to derive greater insight into how changes in chromatin state lead to changes in gene expression, we used the computational package FigR^35^ which connects changes in accessibility of *cis* regulatory elements to changes in gene expression to define domains of regulatory chromatin (DORCs) that are essential in driving the cellular process of interest, in our case neurogenesis.

To identify DORCs essential for neurogenesis, we used our Eupl^WC^ dataset where neurons were being generated appropriately. Applying a cutoff of seven significant chromatin peaks, we identified 34 DORCs including known neurogenic genes such as *roundabout guidance receptor 3* (*ROBO3)*, *neuronal differentiation 4* (*NEUROD4)*, and *contactin 2* (*CNTN2)* (**Fig. 6A**). By clustering changes in accessibility over pseudotime, we identified two clusters of DORCs with distinct accessibility and expression patterns (**Fig. 6B**). One cluster consisting of neuron related DORCs was inaccessible and expressed at low levels at early pseudotime values before becoming accessible and expressed at later developmental pseudotime (**Fig. 6B**). The second cluster consisting of proliferation related DORCs showed the opposite pattern: accessible and expressed at early pseudotime values before becoming inaccessible with lower expression levels at later pseudotime values (**Fig. 6B**). This defines a clear switch between NPC proliferation and neurogenesis in the Eupl^WC^ condition. However, when applying the Eupl^WC^ DORCs to T21^WC^ accessibility and expression along pseudotime, this organization was lost (**Fig. 6B**). As expected, correlations between accessibility and expression for the Eupl^WC^ defined DORCs were significantly weaker in T21^WC^ (**Fig. 6C-E**). We identified a separate set of 40 DORCs driving the T21^WC^ program with only 3 DORCs overlapping between conditions (**Fig. S13A**). There was also no significant difference in correlation strength between the respective Eupl^WC^ and T21^WC^ DORCs (**Fig. S13B**). Analysis of transcription factors either activating or repressing the expression of the DORCs identified 44 transcription factors comprising the gene regulatory network in Eupl^WC^ (**Fig. 6F, Fig. S13C**) while a separate network was identified in T21^WC^ (**Fig. S13D**). For example, the expression of neurogenic DORCs including *ROBO3* (**Fig. 6G**) and *NEUROD4* (**Fig. 6H**) are influenced by multiple transcription factors including *HMGA2* and *PBX homeobox 1* (*PBX1)* both of which showed differential expression between Eupl^WC^ and T21^WC^ NPCs (**Fig. 3C, Fig. S9A**).

**Figure 6.**
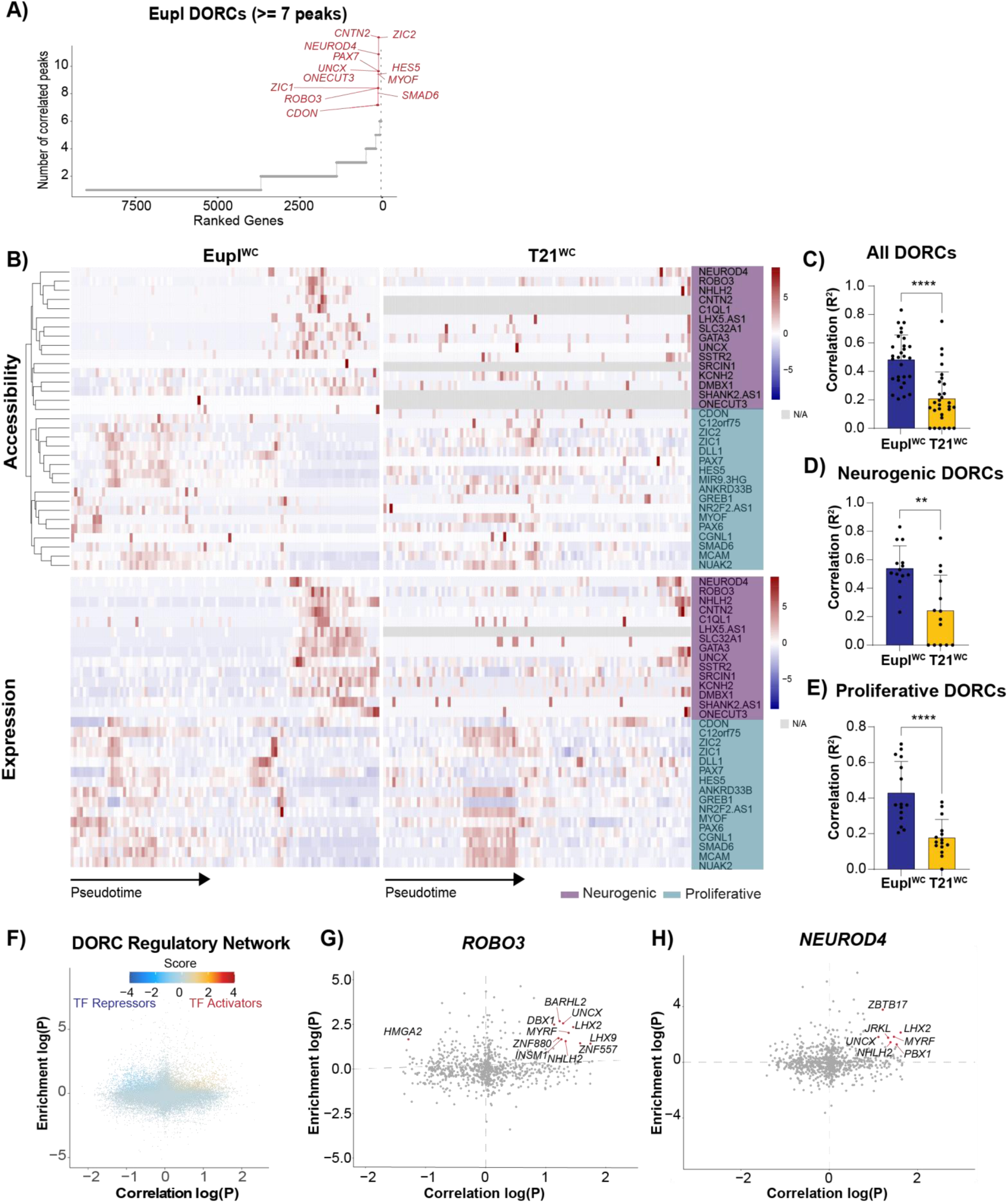
– T21 dysregulates gene regulatory networks driving neurogenesis. **A)** Representation of DORCs identified from the Eupl^WC^ dataset based on a 7 peak cut-off. **B)** Median per cell accessibility and expression level for each Eupl^WC^ defined DORC plotted versus the pseudotime value. Neurogenic DORCs are shown in purple and proliferative DORCs are shown in green. **C-E)** The mean accessibility-expression correlation value for all DORCs (p<0.0001)(C), neurogenic DORCs (p=0.0010)(D), and proliferative DORCs (p<0.0001)(E). Significance was determined by Welch’s t test. **F)** Analysis of transcription factors either activating or repressing the expression of the Eupl^WC^ defined DORCs identified 44 transcription factors (cut-off score>1). **G-H)** Examples of neurogenic DORCs *ROBO3* and *NEUROD4* regulated by transcription factors with differential expression in T21^WC^ NPCs (Fig. 3C, Fig. S7A).

Together, these data indicate that T21^WC^ NPCs failed to follow the molecularly defined Eupl^WC^ neurogenic program. Moreover, the gene regulatory networks driving differentiation were almost entirely separate between T21 and euploid conditions.

### Dysregulated fate-instructive genes in T21 are enriched for heterochromatin marks

Given the prominent cell fate phenotype coupled with changes in chromatin accessibility and broad shifts in molecular program observed in T21, we looked for potential epigenetic signatures using ENCODE data^36^. Strikingly, DEGs identified from each NPC cluster were significantly enriched for heterochromatin marks; the facultative heterochromatin mark histone H3 lysine 27 trimethylation (H3K27me3) was the most significantly enriched histone mark across all five NPC clusters, followed by the constitutive heterochromatin mark histone H3 Lysine 9 trimethylation (H3K9me3), which was significantly enriched across four clusters (**Fig. S14A**). H3K27me3 plays a central role in both the maintenance of pluripotency, and cell fate determination during embryonic development; as a repressive mark deposited by PRC2, H3K27me3 is frequently found at the promoters of developmental genes where it plays a critical role in chromatin condensation, thus restricting transcription factor access and maintaining stemness^37^. During differentiation, proper removal of H3K27me3 at lineage-specific genes is essential to facilitate cell fate transitions^37^. As PRC2 activity can increase chromatin compaction^38^, we also assessed proximal promoters with differentially accessible peaks in T21 NPCs. As with the transcript-level data, H3K27me3 was the most significantly enriched histone mark across all five NPC clusters, followed by H3K9me3 (**Fig. S14B**). Merging all WC NPC clusters and splitting by increased and decreased accessibility revealed that the heterochromatin enrichment was strongest among loci with decreased accessibility in T21 (**Fig. 7A**). Similarly, merging DEGs across all clusters and splitting by up– and down-regulation revealed that the heterochromatin enrichment was strongest among the genes downregulated in T21 NPCs (**Fig. 7A**), consistent with increased gene repression.

**Figure 7.**
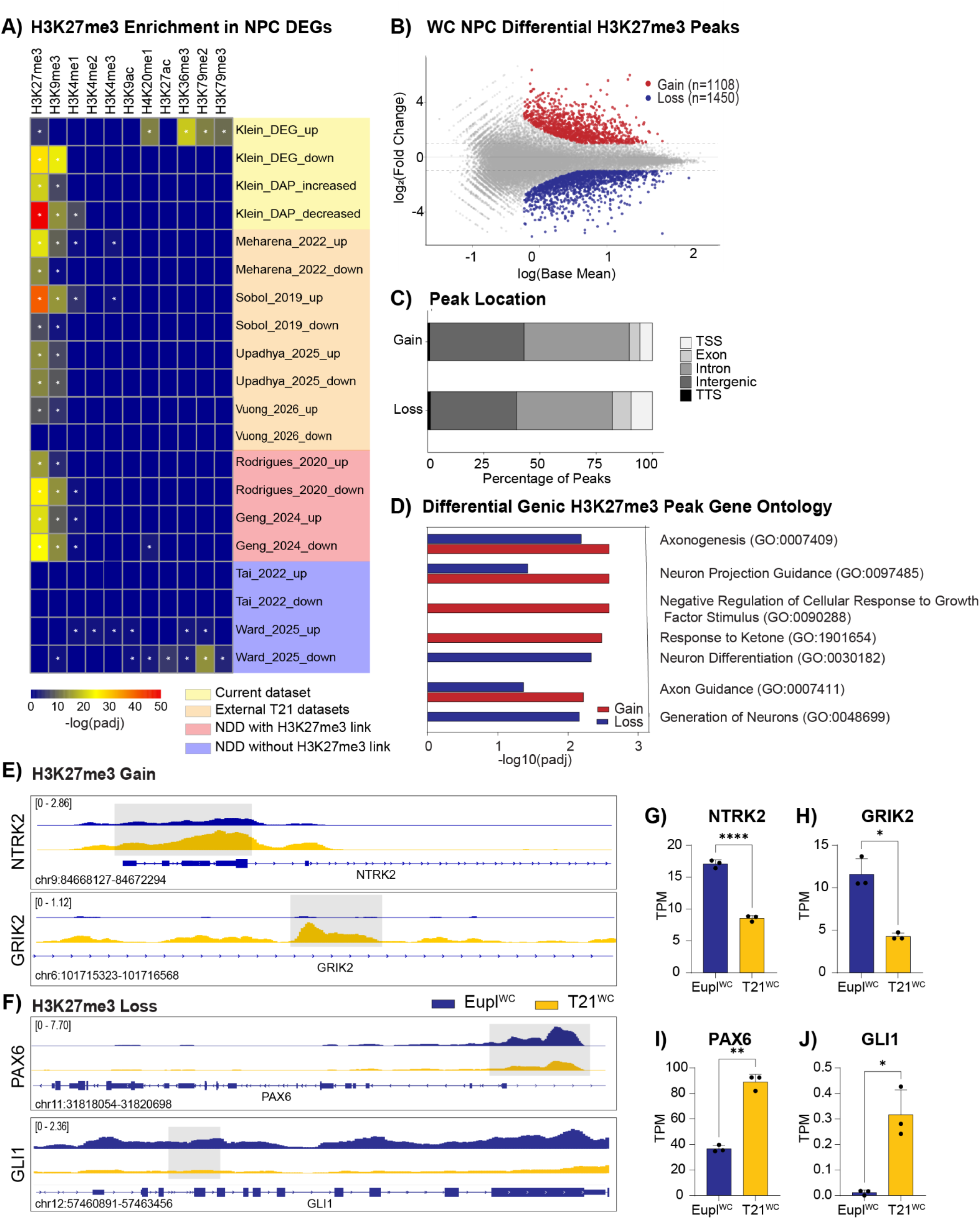
– Redistribution of H3K27me3 binding in T21 NPCs. **A)** Comparison of significantly upregulated or downregulated DEGs from combined NPC clusters (Klein; this study), four additional published T21 datasets (Meharena et al 2022; Sobol et al 2019; Upadhya et al 2025; Vuong et al. 2026), two transcriptional datasets from mutations expected to impact heterochromatin (Rodrigues et al 2020; Geng et al 2024), and two datasets from 16p11.2 duplication (Tai et al 2022; Ward et al 2025) with the ENCODE Histone Modifications 2015 database, * indicates p-value < 0.05. **B)** Differential H3K27me3 peaks are both gained (n=1108) and lost (n=1450) in T21^WC^ NPCs (l2FC>1; p-value <0.05). **C)** Genomic location of gained and lost H3K27me3 peaks. **D)** Top five significant gene ontology terms of the gained and lost peaks. E) Example gene tracks where T21^WC^ NPCs gained H3K27me3 peaks and F) example gene tracks where T21^WC^ NPCs lost H3K27me3 peaks compared to Eupl^WC^ NPCs. G) *NTRK2* (p<0.001) and **H)** *GRIK2* (p=0.0168) show a significant decrease in expression in T21^WC^ NPCs and **I)** *PAX6* (p=0.0013) and **J)** *GLI1* (p=0.0315) show a significant increase in expression in T21^WC^ NPCs compared to Eupl^WC^ NPCs. Significance was determined by Welch’s t test. A minimum of n=3 independent wells per condition per karyotype were used for each experiment. Each well was independently passaged and differentiated starting from the iPSC stage to generate biological replicates.

To assess the generalizability of these results, we analyzed the histone enrichment patterns from three published T21 NPC transcriptional datasets and one fetal brain dataset. In all datasets queried, H3K27me3 was the top enriched histone mark (**Fig. 7A**). Notably, the levels of H3K27me3 and H3K9me3 enrichment observed for genes dysregulated in human T21 NPCs was equivalent to the levels of enrichment in human NPCs harboring mutations with explicit links to heterochromatin. Genes dysregulated by loss of AUTS2^39^, which can interact with PRC2^40^, as well as genes dysregulated by mutations in MECP2^41^ which can interact with H3K27me3^42,43^, showed levels of H3K27me3 and H3K9me3 enrichment equivalent to the T21 datasets (**Fig 7A**). By contrast, no heterochromatin enrichment was observed for genes dysregulated by 16p11.2 duplication^44,45^ (**Fig 7A**). Moreover, given the cell fate changes previously identified in the hematopoietic system in T21, we also examined four published blood cell datasets^20,21,46,47^; here, histone enrichment was only present in the downregulated genes, with H3K27me3 again the top enriched mark across all datasets (**Fig. S14C**). Collectively, these data unexpectedly identify a common thread of heterochromatin dysregulation across T21 progenitor cells from diverse donor and tissue sources.

### Genome-wide redistribution of H3K27me3 in T21 NPCs

To further investigate the H3K27me3 landscape in T21, we performed Cleavage Under Targets and Tagmentation (CUT&Tag) on Eupl^WC^ and T21^WC^ P3 NPCs (**Fig. 7B-J; Fig. S15**). We observed a broad redistribution of H3K27me3 with roughly the same number of peaks gained (n=1108) and lost (n=1450) in T21 NPCs compared with isogenic euploid control NPCs (**Fig. 7B; Table S4**). The differential peaks were distributed across the genome, with a majority in non-coding regions (**Fig. 7C**). For differential peaks in coding regions, ontology analysis pointed to an altered regulatory landscape for genes driving neurogenic processes with terms such as Axogenesis, Neuron Projection Guidance, and Neuron Differentiation significantly overrepresented among genes with differential H3K27me3 binding (**Fig. 7D**). More specific examination of the genes contained within these terms identified increased H3K27me3 binding on genes that drive neurogenesis such as *Neurotrophic receptor tyrosine kinase 2 (NTRK2)* and *Glutamate Ionotropic Receptor Kainate Type Subunit 2 (GRIK2)* (**Fig. 7E**) along with corresponding decreases in transcript level (**Fig. 7G-H**) while genes associated with NPC fate and cycling such as *PAX6* and *GLI family zinc finger 1 (GLI1)* had decreased H3K27me3 binding (**Fig. 7F**) and corresponding increases in transcript level in T21 NPCs (**Fig. 7I-J**). These results are consistent with altered H3K27me3 landscapes influencing differentiation potential in T21 NPCs.

### EZH2 inhibition increases neurogenesis in T21

Given the significant changes to H3K27me3 binding patterns observed in T21, we assessed the functional impacts of PRC2 inhibition on NPC fate change in T21. Several recent studies have successfully employed PRC2 inhibition to increase NPC differentiation in the non-diseased state^48,49^. Specifically, we used pharmacological inhibition of enhancer of zeste homolog 2 (EZH2i), a catalytic component of PRC2 on T21^WC^/Eupl^WC^ NPCs at multiple time-points over the course of differentiation compared to vehicle control (**Fig.8A; Fig. S16A-F**). NPCs were treated with DMSO or 2µM of the EZH2i GSK126 for all four weeks of NPC induction, for the last three weeks of NPC induction, or for the last two weeks of NPC induction, followed by spontaneous differentiation (**Fig. 8A**). As expected, T21^WC^ NPCs treated with EZH2i showed a significant reduction in levels of H3K27me3 and gain of the active mark histone H3 lysine 27 acetylation (H3K27ac) after two, three, or four weeks of treatment compared to vehicle control (**Fig. 8B-C, Fig. S16A-B**).

**Figure 8.**
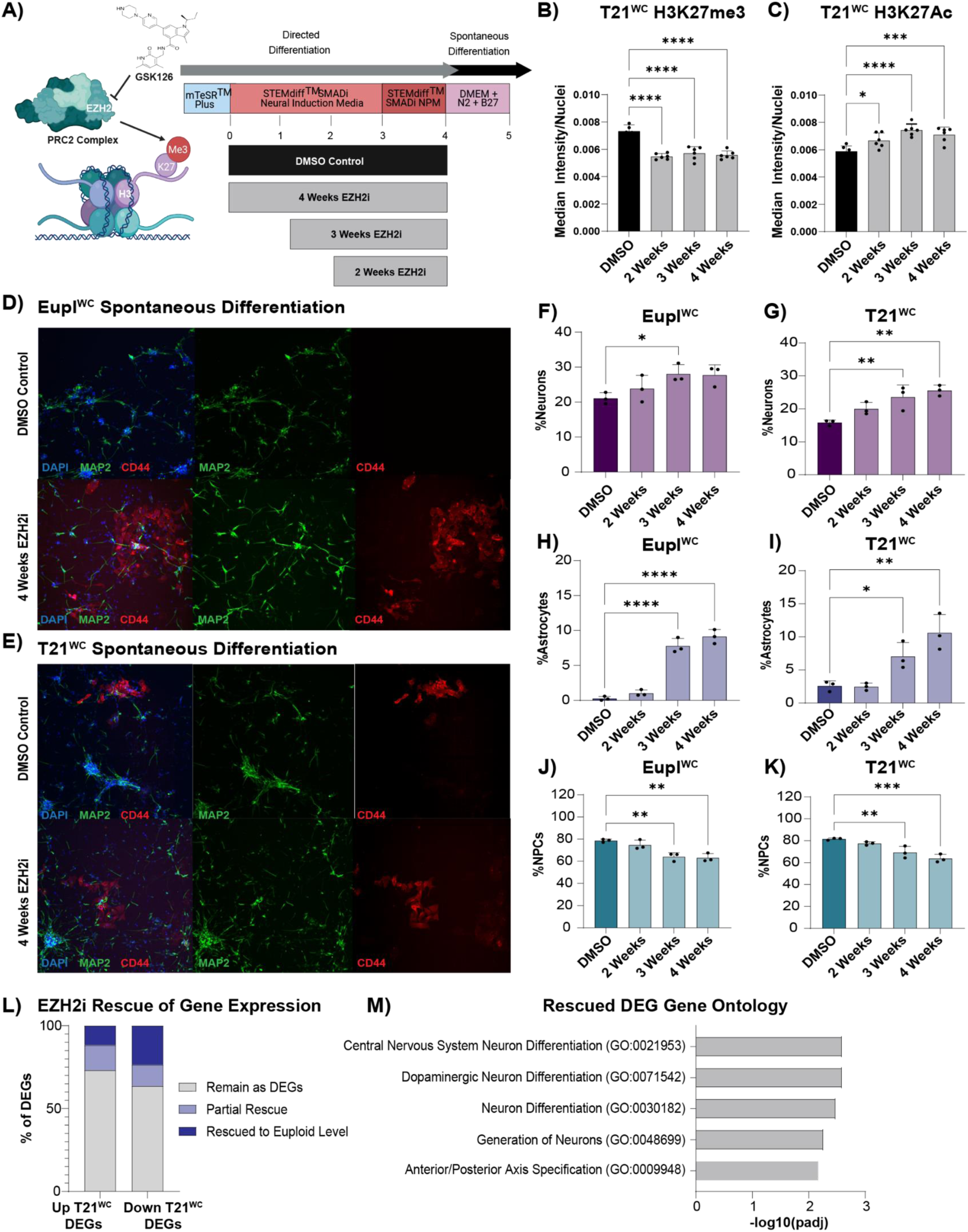
– EZH2 inhibition increases neurogenesis in T21. **A)** Schematic of EZH2i treatment paradigm (Biorender). **B)** Quantification of H3K27me3 levels (median intensity/nuclei) in T21^WC^ NPCs treated with EZH2i (2 weeks p<0.0001; 3 weeks p<0.0001; 4 weeks p<0.0001). **C)** Quantification of H3K27ac levels (median intensity/nuclei) in T21^WC^ NPCs treated with EZH2i (2 weeks p=0.0271; 3 weeks p<0.0001; 4 weeks p=0.0009). **D-E)** Representative immunocytochemistry of the Eupl^WC^ (D) and T21^WC^ (E) cells post spontaneous differentiation after either DMSO or 4 weeks of EZH2i treatment. Cells are stained with MAP2 (neurons) and CD44 (astrocyte precursors) and counterstained with DAPI. **F)** Eupl^WC^ cells treated for three weeks with EZH2i produced significantly more neurons than DMSO treated controls (28.1% ± 2.6% 3 weeks vs 21.1% ± 1.6% DMSO p=0.0423). **G)** T21^WC^ cells treated for three weeks and four weeks with EZH2i produced significantly more neurons than DMSO treated controls (23.6 ± 3.7% 3 weeks and 25.6% ±1.6% 4 weeks vs 15.8% DMSO p=0.0077 and p=0.0019 respectively). **H)** Eupl^WC^ cells treated for three weeks and four weeks with EZH2i produced significantly more astrocyte precursors than DMSO treated controls (7.8% ± 1.1% 3 weeks and 9.1% ± 1.0% 4 weeks vs 0.3% ± 0.3% DMSO p<0.0001 and p<0.0001 respectively). **I)** T21^WC^ cells treated for three weeks and four weeks with EZH2i produced significantly more astrocyte precursors than DMSO treated controls (7.1% ± 2.1% 3 weeks and 10.6% ± 2.7% 4 weeks vs 2.6% ± 0.8% DMSO p=0.0377 and p=0.0014 respectively). **J)** Eupl^WC^ cells treated for three weeks and four weeks with EZH2i produced significantly fewer NPCs than DMSO treated controls (64.1% ± 3.7% 3 weeks and 63.1% ± 3.8% 4 weeks vs 78.6% ± 1.4% DMSO p=0.0025 and p=0.0016 respectively). **K)** T21^WC^ cells treated for three weeks and four weeks with EZH2i produced significantly fewer NPCs than DMSO treated controls (69.4% ± 5.5% 3 weeks and 63.8% ± 3.7% 4 weeks vs 81.6% ± 0.9% DMSO p=0.0063 and p=0.0006 respectively). Significance was determined by One-way ANOVA followed by Dunnett tests for multiple comparisons. **L)** The expression level of DEGs identified between T21^WC^ and Eupl^WC^ DMSO NPCs were either partially or fully restored to euploid levels after four weeks of EZH2i (Up DEGS: 15.2% partially and 11.6% fully rescued. Down DEGs: 12.9% partially and 23.5% fully rescued). M) Gene ontology of the fully and partially rescued DEGs post EZH2i treatment. A minimum of n=3 independent wells per condition per karyotype were used for each experiment. Each well was independently passaged and differentiated starting from the iPSC stage to generate biological replicates.

To assess the impact of H3K27me3 loss on neural cell fate, cells from each karyotype were first assessed for proportions of neurons, astrocyte precursors, and NPCs following spontaneous differentiation. While no changes were observed after two weeks of EZH2i, three or more weeks of EZH2i drove significant increases in the percentage of neurons and astrocyte precursor cells produced by both karyotypes alongside a concurrent decrease in percentage of NPCs consistent with enhanced differentiation (**Fig. 8D-K**). While the percentage of differentiated cells increased in each karyotype, the magnitudes of effect differed. Notably, the percentage of neurons in the Eupl^WC^ condition increased by 33.3% (three week EZH2i) and 31.6% (four week EZH2i) compared to control, while the percentage of neurons in the T21^WC^ condition increased by 48.9% (three week EZH2i) and 61.6% (four week EZH2i) compared to control (**Fig. 8D-G**). Thus, after 3-4 weeks of EZH2i, T21^WC^ and Eupl^WC^ achieved similar proportions of cell types (**Fig. 8D-K**). These data are consistent with previous studies showing enhanced NPC differentiation only after several weeks of PRC2 inhibition^48,49^. Moreover, global transcriptional analyses of NPCs after four weeks of EZH2i versus DMSO control revealed that 14.6% of DEGs in T21 were fully restored to euploid expression levels with EZH2i and another 14.8% moved in the direction of euploid expression (**Fig. 8L, Table S4**). As expected, EZH2i had a larger impact on restoring the expression levels of genes that were downregulated in T21 compared to genes that were upregulated in T21, with 36.4% of the downregulated DEGs showing at least a partial rescue compared to 26.8% of the upregulated genes (**Fig. 8L, Table S4**). Consistent with the immunophenotyping data, GO analysis of the restored genes revealed significant enrichment for neuron development terms such as Central Nervous System Neuron Differentiation and Generation of Neurons (**Fig. 8M**). Together, these data provide proof-of-principle that PRC2 inhibition is sufficient to specifically overcome the early neurogenesis block in T21.

## DISCUSSION

Our results using *in vitro* human models recapitulate the prominent cell fate phenotypes observed in T21. Through rigorous cell cycle analyses, we find a significant shift in the G1/S ratio evident from the earliest time-point of NPC induction, with T21 cells spending more time in S phase and less time in G1, a phenotype consistent with continued proliferation rather than differentiation. At this stage, neurogenic gene expression programs were significantly decreased in T21 with a concomitant decrease in chromatin accessibility at proximal promoters of neurogenic genes, and enrichment for heterochromatin marks. The failure of T21 NPCs to activate an orchestrated neurogenic program at the chromatin level, coupled with the lack of evidence for premature neuronal differentiation, a strong astroglial driver signature, or increased neuronal cell death collectively support the hypothesis that T21 NPCs inappropriately maintain a closed chromatin structure over neurogenic loci, leading to reduced neurogenesis during early stages of differentiation. Moreover, genome-wide re-wiring of H3K27me3 landscapes in T21 combined with the impacts of pharmacological EZH2i support the hypothesis that aberrant chromatin remodeling contributes to neural cell fate changes in T21.

As discussed above, one previous study using human fetal tissue reported dynamic changes in progenitor populations in T21 including more basal and intermediate progenitor cells at early to mid time-points, followed by a decrease in these populations at later timepoints^11^. We speculate that our *in vitro* models, which reflect very early stages of neurogenesis, could be capturing a window of aberrant NPC expansion in T21, which may be followed by a premature decrease in T21 NPCs at later developmental stages. This would be consistent with the increased number of NPCs that we observed in T21 across multiple independent assays, as well as the reduced neuron output. While we did not detect increased senescence or cell death in T21 using P3 NPCs, it’s also possible that remaining NPCs senesce or die at later timepoints, which would be consistent with studies reporting the emergence of a senescence phenotype in T21 at later developmental stages^50^. The intrinsic differences in proliferation rates that we identified between cell lines also provided an opportunity to assess correlations between overall proliferation rate and neuronal output. Here, even T21 cells with faster cycling maintained decreased neuronal production, suggesting that differences in proliferation rates were not sufficient to account for the neurogenesis phenotype. The G1/S shift we identified in T21, which was independent of overall proliferation rate, more likely reflects a decrease in neurogenic versus proliferative divisions between karyotypes, consistent with studies linking G1/S ratios and neurogenesis in other cellular systems^27–29^.

We leveraged PRC2 inhibition to provide proof-of-principle that decreased neurogenesis is driven in part through a chromatin mechanism, but it remains to be determined how longer-term PRC2 inhibition or treatment at different stages of corticogenesis would impact final cellular output. Multiple studies have examined the role of H3K27me3 in the neurogenic to astrogliogenic fate switch. For example, studies in mouse cortical NPCs^51^ and human cortical organoid models^48^ report that loss of PRC2 can promote NPC differentiation, reducing the neurogenic window and leading to a reduction in neuronal output. By contrast, PRC2 has been shown to restrict neurogenesis in mouse cortical NPCs and promote astrogliogenesis, with loss of PRC2 components leading to prolonged neurogenesis and delayed fate switching^52^. Consistent with this latter finding, human mutations in a H3K27 demethylase (*KDM6A*; leading to increased H3K27me3) are frequently associated with microcephaly, while mutations in core PRC2 components (*EZH2*; leading to reduced H3K27me3) are frequently associated with macrocephaly^53^. In mouse models, PRC2 has also been reported to function in a non-cell autonomous manner in neurogenesis, while it cell autonomously regulates astrocyte proliferation and maturation^54^ consistent with distinct mechanisms of PRC2 at different steps of corticogenesis. Importantly, our data as well as the above studies all strongly support a role for PRC2/H3K27me3 in the timing of NPC fate switching during cortical development while underscoring a more complex picture which may include both cell intrinsic and cell extrinsic mechanisms coupled with regional/cell type diversity, which likely contribute to diverse outcomes.

T21 is characterized by significant inter– and intra-individual variation^55,56^ despite individuals sharing the same chromosomal alteration. Our data support shared dysregulation of heterochromatin marks in NPCs from diverse donors and tissue types, despite variability in expression changes. Indeed, all nine transcriptional datasets from human T21 progenitor cell types queried showed significant enrichment for heterochromatin marks. Our results are also generally consistent with a previous study showing chromatin remodeling during fetal hematopoiesis in T21 driving changes in erythroid lineage specification^57^. There, multiomic data from blood cell types suggested that T21 can “prime” hematopoietic stem cells to differentiate toward the erythroid lineage, with enhanced correlation between gene expression and chromatin accessibility^57^, which was not present in our NPC data. We speculate this may reflect cell type, cell state, or genetic background differences. Notably, there are many potential modifiers which could differentially impact H3K27me3 or H3K9me3 binding at individual loci, including differences in the expression of both *cis* and *trans* regulatory elements. Overall, our data are consistent with heterochromatin dysregulation driving early neurodevelopmental defects in T21. This also provides a potential explanation for how shared phenotypes in T21 (i.e., reduced neurogenesis) could be driven through diverse gene expression changes, depending on heterochromatin formation at specific loci which can be impacted by multiple genetic and environmental factors. Further studies will be required to assess specific patterns of dysregulation across donors and tissue types.

Ultimately, the mechanisms driving chromatin compaction at neurogenic loci remain to be determined. It is possible that increased dosage of individual genes encoded on HSA21 drives phenotypes, including epigenetic factors (i.e., *BRWD1, HMGN1, DNMT3L*). Although none have direct links to maintenance of a repressive chromatin structure, overexpression of the HSA21-encoded gene HMGN1 in B cells has been linked to H3K27me3 repression^58^. It’s also important to note that in studies from yeast to mammals, altered gene-dosage has been shown to drive defects in proliferation, metabolism, and proteostasis, regardless of the identity of the extra chromosome and consistent with a general cellular response to aneuploidy^59–64^. Recent studies highlight the key roles that diverse metabolic pathways play in epigenetic regulation during cortical development, with molecules such as alpha-ketoglutarate and S-adenosylmethionine key for histone methylation^53^. Links between T21 and altered metabolism are well-established^65–67^ and it is possible that metabolic changes in T21 influence histone modifications. Chromatin accessibility and histone modifications are also downstream of higher-order topological features (i.e., A/B compartments, LADs, TADs, enhancer-promoter loops, chromatin compartments) which can alter the repressive chromatin environment. While limited studies have examined the effects of large-scale CNVs on the 3D genome architecture, those which have, report significant genome-wide findings^68,69^. It will be particularly interesting to explore in future studies whether metabolic dysregulation or disruption of 3D genome architecture could specifically contribute to chromatin changes driving cell fate change in T21. Alternatively, there could be multiple, independent routes to heterochromatin dysregulation, reduced neurogenesis, and ultimately brain undergrowth^54^. Disentangling these possibilities is essential to understand how diverse alterations including aneuploidies negatively impact human neural development.

This study has several limitations. Our results focus on NPC fate specification but do not explore differences in the identity or maturation state of neurons or astrocyte precursors, and the relationship between these phenotypes and chromatin structure. This is in part due to the lack of sufficient mature cell types generated from both karyotypes for powered comparisons. With a significant increase in scale, it would be interesting to examine gene expression and chromatin accessibility profiles across post-mitotic neural cell types. Based on studies of fetal and post-mortem tissue, we might expect to observe differences in neural subtype identities between karyotypes, such as a shift in the ratio of upper layer to deep layer neurons^11^. Neuron generation through dual-SMAD inhibition also tends to drive the generation of neurons with a deep layer fate so it will be important to consider additional time-lines and paradigms to capture both upper and deep layer phenotypes. Moreover, analyses of third trimester human fetal development have revealed that the timing of the neurogenic to astrogliogenic fate switch varies by cortical region^70^. Regional differences in cortical expansion *in vivo* are noteworthy as they suggest that additional cell intrinsic (e.g., specific NPC subtypes) or cell extrinsic (e.g., microenvironmental niche) factors influence neurogenesis, underscoring the need for additional datasets from diverse NPC subtypes or brain regions. Additionally, while we analyze independent isogenic iPSC pairs, cellular resources linked to patient data (e.g., degree of cognitive impairment, presence of co-occurring conditions, brain imaging, etc.) are missing from the field and required to facilitate deeper investigation of genotype-phenotype relationships.

Collectively, this study provides an in-depth analysis of early neural cell fate change in T21, connecting Polycomb dysregulation to this key phenotype, thus enhancing insight into the intricate mechanisms underlying human development and their dysregulation in autosomal aneuploidy.

## METHODS

### Cell culture

Two isogenic human iPSC line pairs discordant for trisomy 21 were purchased from WiCell and used in experiments as noted. The female line consisted of WC-24-02-DS-B (Eupl^WC^) and WC-24-02-DS-M (T21^WC^) while the male lines consisted of UWWC1-DS2U (Eupl^DS^) and UWWC1-DS1 (T21^DS^). Both isogenic lines were maintained in MTeSR Plus media (STEMCELL Technologies, 100-0276) supplemented with penicillin/streptomycin and Normocin (InvivoGen, ant-nr-05) on Geltrex (ThermoFisher, A14132-02) coated plates and passaged with TrypLE (ThermoFisher) plus ROCKi. For NPC differentiation, iPSCs were induced using the STEMDiff SMADi Neural Induction Kit (STEMCELL Technologies, 08581). NPCs were seeded at a density of 2×10^6^ cells per well in a 6-well plate, with 3 wells per karyotype. Media was changed daily, and Accutase (VWR, 490007-741) was used to passage cells weekly following the initial induction. After three weeks, NPCs were transferred to StemDiff Neural Progenitor Media (STEMCELL Technologies, 05833) for one additional week. At the end of this passage cells were considered NPCs based on SOX1 expression and morphology and were either collected for downstream analysis or maintained in StemDiff Neural Progenitor Media. For spontaneous differentiation, P3 NPCs were passaged and plated at a density of 0.4×10^6^ cells per well in a 6 well plate. Cells were cultured in DMEM/F12 (ThermoFisher, 11320033) supplemented with 1x N-2 (Gibco, 17502048) and 1x B27 (ThermoFisher, 17504044-044). After one week of spontaneous differentiation, cells were either collected or fixed for downstream analysis. All cells were grown in a 5% CO2 and 37°C incubator and routinely tested for mycoplasma contamination. G-banding karyotype analysis was used to confirm expected karyotypes (Cell Line Genetics). A minimum of n=3 independent wells per condition per karyotype were used for each experiment. Each well was independently passaged and differentiated starting from the iPSC stage to generate biological replicates.

### GFP clonal analyses

Eupl^WC^ and T21^WC^ iPSCs were dually infected with ultra-high titer Tet-O-GFP and rtTA lentivirus (Alstem) to produce stable lines of cells with doxycycline inducible GFP expression. iPSCs were karyotyped (Cell Line Genetics) to ensure lentiviral infection did not result in chromosomal abnormalities. Infected iPSCs were differentiated into NPCs using the method described above. At passage 2, the GFP labeled NPCs were mixed with non-infected Eupl^WC^ and T21^WC^ NPCs of the same karyotype at a ratio of 1:10,000. The mixed cultures were passaged and allowed to spontaneously differentiate as described above.

### EZH2 inhibition

T21^WC^ and Eupl^WC^ NPCs were treated daily with 2 µM GSK126 (Selleckchem, S7061) resuspended in DMSO or an equivalent volume of DMSO as vehicle control. Briefly, for NPCs treated for four weeks, daily additions of GSK126 to the Neural Induction Media began the day after NPC induction, for NPCs treated for three weeks, the daily additions started the day of the first NPC passage, and for NPCs treated for two weeks, the daily additions started the day of the second NPC passage. Drug treatment continued until the NPCs were plated for spontaneous differentiation as described above.

### Immunocytochemistry and EdU staining

Cells were washed with 1x PBS and fixed for 20 minutes in 4% paraformaldehyde plus 4% sucrose in 1x PBS. Fixed cells were washed in 1x PBS and stored at 4°C for up to one week prior to staining. Fixed cells were permeabilized and blocked for 20 minutes in 4% Horse serum (ThermoFisher, 16050114), 0.3% Triton X-100 (Millipore Sigma, T9284) and 0.1M Glycine (Millipore Sigma, G7126) in 1x PBS at room temperature. This was followed by incubation of the primary antibodies in 4% Horse serum in 1x PBS at 4°C overnight. After incubation, the cells were washed with 0.3% Triton X-100 in 1x PBS then incubated with the secondary antibodies along with DAPI (1:5000, ThermoFisher, D1306) and TrueBlack (1:5000, 568 Biotium, 23007) in 4% Horse serum in 1x PBS for one hour at room temperature. Finally, cells were abundantly washed with 1x PBS and stored at 4°C prior to imaging. The following primary antibodies were used: rat anti-CD44 (1:500, ThermoFisher, 14-0441-82), chicken anti-MAP2 (1:1000, Abcam, ab5392), goat anti-SOX1, (1:500, R&D Systems, AF3369), rabbit anti-GFP (1:500, Abcam, ab6556), rabbit anti-CC3 (1:500, Cell Signaling Technologies 9661), rabbit anti-pINK4a (1:500, ThermoFisher MA5-32133, mouse anti-Ki67 (1:1000, Cell Signaling Technologies 9449), rabbit anti-PH3 (1:500, Millipore Sigma 06-750), rabbit anti-H3K27me3 (1:2000, Active Motif, 91167), and rabbit anti-H3K27ac (1:2000, Cell Signaling, 8173T). The following AlexaFluor secondary antibodies were used: goat anti-rat 647 (A21247), donkey anti-chicken 488 (A78948), goat anti-chicken 555 (A21437), donkey anti-goat 647 (A21447), donkey anti-rabbit 488 (A21206), donkey anti-mouse 647, and donkey anti-rabbit 555 (all 1:2000 dilution, ThermoFisher). To mark cells in S-phase, we used the EdU Assay / EdU Staining Proliferation Kit (Abcam, ab219801). NPCs were incubated with 10 µM EdU for two hours then fixed and processed for imaging according to the kit protocol.

### Image acquisition and analysis

For bulk phenotyping and proliferation assays, post immunostaining, images were captured using the Opera Phenix high content screening system (Perkin Elmer). Spontaneous differentiation experiments were acquired using a 10x objective lens to capture a larger field, while proliferation stains were captured with a 20x objective lens. For each condition, 5-10 fields were chosen and imaged at random in 3-6 wells resulting in a total of 30 images per replicate. Images were analyzed using CellProfiler (version 4.2.8) to obtain cell counts or stain intensity within the outlined nucleus for each stain. Outliers were identified and removed using GraphPad Prism. Counts for each replicate were averaged and statistical analysis was performed as described in the text using GraphPad Prism. For clonal analyses, images were captured using a Lionheart FX Automated Microscope (Agilent) to allow targeted capture of individual GFP expressing cells and progeny. A minimum of 20 clones were captured per condition. Images were analyzed using ImageJ (version 1.53q) and statistical analysis was performed using GraphPad Prism.

### Flow Cytometry

Each condition was plated in triplicate in a 6-well dish with each sample treated individually. Cells were collected half-way through their noted passage time to ensure their growth was not constrained by density. Cells were treated with Accutase and filtered to form a single cell suspension. One million cells per replicate were incubated with Vybrant DyeCycle Green Stain (ThermoFisher, V35004) at 37°C for 30 minutes, then incubated with SYTOX Red Dead Cell Stain (ThermoFisher, S34859) at 4°C for 15 minutes. Cells were analyzed at the Harvard Stem Cell and Regenerative Biology Flow Cytometry Core on a BD FACSAria II. 50,000 cells per sample were analyzed. FlowJo (Version 10.10.0) was used for sample analysis. Samples were first gated based on size to remove debris and doublets. Dead cells based on SYTOX Red positivity were removed from analysis and the remaining VYBRANT Green positive cells were used to determine cell cycle phase. The Watson (pragmatic) model was used to determine the proportion of cells in G1, S, and G2. The model was constrained so that G1=G2 and the optimum peak range was determined for each sample based on the minimal Root Mean Square Deviation value. To assess changes in proliferation rate, each condition was plated in triplicate in a 6-well dish and treated individually. At the beginning of each passage noted, cells were incubated with 5 µM CellTrace Violet (ThermoFisher, C34557) in DMEM/F12 (ThermoFisher, 11320033) at room temperature for seven minutes then processed according to the kit protocol. After 48 hours of culture, cells were collected for analysis at the Broad Technology Space on a CytoFLEX LX Flow Cytometer (Beckman Coulter). For each karyotype the mean CellTrace Violet intensity was compared between cells cultured for 48 hours and cells freshly stained. The change in intensity was compared between karyotypes to produce a relative proliferation rate for each condition.

### TMT-Proteome

Frozen cell pellets were lysed by the addition of 500 uL of lysis buffer (75 mM sodium chloride, 10 mM sodium pyrophosphate, 10 mM sodium fluoride, 10 mM β-Glycerophosphate, 10 mM sodium orthovanadate, 50 mM EPPS pH 8.5, cOmplete^TM^ Mini EDTA-free Protease Inhibitor Cocktail (Sigma Aldrich), 3 % SDS, 5 mM PMSF). Disulfide bonds were reduced with dithiothreitol (DTT) at a final concentration of 5 mM for 30 minutes at 56 °C, and then alkylated with iodoacetamide (IAA) at a final concentration of 15 mM for 20 min at room temperature in the dark. The reaction was quenched with DTT (5 mM final concentration) for 20 minutes at room temperature in the dark. Protein was reduced, alkylated, purified through TCA precipitation^71^, and digested with Lys-C (Wako Chemicals) and trypsin (Promega) as described previously^72^. Peptide concentrations of each sample were determined using a BCA assay (Thermo Fisher Scientific). Peptides were labeled using TMTpro^73^ reagents (Thermo Fisher Scientific) by resuspending 25 µg of peptides in 25 µL of 200 mM EPPS (pH 8.5), 30% acetonitrile (ACN), and adding 325 µg TMT reagent from a stock in anhydrous ACN. Upon an hour at room temperature, the reaction was quenched with hydroxylamine (0.5 % final concentration). The labeled digests were acidified with TFA (1 % final concentration), pooled, and desalted as described previously^72^.

The pooled samples were prefractionated using basic pH reversed-phase liquid chromatography (bRPLC)^72^ and analyzed by LC-M2/MS3 on an Orbitrap Fusion Lumos mass spectrometer using the Simultaneous Precursor Selection (SPS) supported MS3 method^74,75^ as described previously^76^. MS2 spectra were assigned using a SEQUEST-based^77^ in-house built proteomics analysis platform^78^ using a target-decoy database-based search strategy to assist filtering for a false-discovery rate (FDR) of peptide and protein identifications of less than 1 %^78,79^. Peptides matching multiple protein sequences were assigned to the protein explaining the largest number of matched redundant peptide sequences following the law of parsimony^78^. TMT reporter ion intensities were extracted from the MS3 spectra, selecting the most intense ion within a 0.003 m/z window centered at the predicted m/z and spectra were used for quantification if the average signal-to-noise ratio per channel exceeded 20 and the isolation specificity^74^ for the precursor ion was ≥ 0.75. Protein intensities were calculated by summing the TMT reporter ions for all peptides assigned to a protein. Intensities were first normalized by the average intensity across all samples relative to the median average across all proteins^76^ followed by normalization of protein intensities determined for each sample by the average of the median intensities across the samples^76^.

### Total RNAseq

Triplicate samples of P3 NPCs were collected following four weeks of either DMSO or GSK126 treatment. Total RNA was isolated and DNase treated using the RNeasy Plus Kit (Qiagen) and following the manufacturer’s protocol. Depletion of rRNA was performed with QIAseq FastSelect rRNA HMR reagents as per the manufacturer’s instructions. Libraries were generated using Roche Kapa Biosystems RNA HyperPrep reagents with a Beckman Coulter Biomek i7. Following pooling of uniquely dual indexed libraries, sequencing was performed with paired-end 150bp reads on a NextSeq 300. Library preparation and sequencing was performed by the Dana-Farber Cancer Institute Molecular Biology Core Facility.

### 10x Genomics sample preparation and sequencing

Three biological replicates of either P3 NPCs or spontaneously differentiated cells per karyotype were disassociated with Accutase and pooled together in a single cell suspension. One million cells per karyotype were collected for 10X Genomics Single Cell Multiome ATAC + Gene Expression prep. Nuclei were isolated following the protocol from Illumina and approximately 8,000 nuclei per condition were submitted for library preparation and sequencing to the Bauer Core Facility at Harvard University. The cDNA library was made using the 10X Genomics Chromium Single Cell 3′ Library and Bead Kit v3. Sequencing was done on NovaSeq_S4. The P3 T21 sample had 1,213 median genes per cell and 19,814 median high-quality fragments per cell, while the spontaneously differentiated T21 sample had 2,661 median genes per cell and 29,798 median high-quality fragments per cell. The P3 euploid sample had 1,089 median genes per cell and 17,838 median high-quality fragments per cell, while the spontaneously differentiated euploid sample had 1,042 median genes per cell and 16,994 median high-quality fragments per cell. Paired-end read Fastq files were aligned to GRCh38 Genome Reference Consortium Human Reference 38 (hg38) using the ‘count’ method of CellRanger (version 8.0.1.4)^80^.

### Single nuclei bioinformatic analyses

#### snRNA-seq QC, integration, normalization, and clustering

snRNA-seq analysis was performed using the R package Seurat (version 5.3.0)^81^. Cutoffs for UMI count, percent mitochondrial genes, and number of unique expressed genes were employed to subset out low-quality cells from each sample. SCTransform was used to normalize each sample individually. DoubletFinder was performed where non-aggregated data was retrieved^82^. The pK parameter was set to the maximum of the bimodality coefficient distribution. The homotypic doublet proportion estimate was calculated using the 10X Genomics multiplet rate per cells recovered (Chromium Single Cell V(D)J Reagent Kits with Feature Barcoding technology for Cell Surface Protein, Document Number CG000186 Rev A, 10x Genomics, (2019, July 25)). Multiplet rates of 5% and 2% were utilized for the P3 and spontaneously differentiated euploid samples and 4.5% and 2% for the P3 and spontaneously differentiated T21 samples. All samples were then merged. The CellCycleScoring function was used to annotate cell-cycle effects, where the difference between the G2M and S phase scores were regressed out in a subsequent call for SCTransform. Principal component analysis (PCA) was done on the Pearson residuals output from SCTransform. Data were then integrated using the Harmony method^81,83^. The top 30 components were then used for uniform manifold approximation and projection (UMAP). Clusters were then identified with FindNeighbors and FindClusters, with the resolution parameter for FindClusters method set to 0.175. Clusters were identified and named based on expression of canonical cell markers.

#### snATAC QC, integration, normalization

snATAC-seq analysis was performed using the R package Signac (version 1.15.0)^84^. In addition to the cutoffs for UMI count, percent mitochondrial genes, and number of unique expressed genes, additional cutoffs for number of total peaks, enrichment within transcription start sites, fraction of reads in peaks, nucleosome banding pattern, and ratio of reads in genomic blacklist regions, or regions associated in having artefactual signal were implemented to subset out low-quality cells of both snRNA-seq and snATAC-seq data. After QC, subsequent steps till merging the four samples follow that of the snRNA-seq pipeline. After merging, snATAC-seq data was normalized using term frequency-inverse document frequency (TF-IDF) normalization with the Signac method RunTFIDF. Top features were identified with the FindTopFeatures for dimensional reduction. Then, the Signac method runSVD performed the singular value decomposition calculation for dimensionality reduction. The combination of TF-IDF and SVD is referred to as latent semantic indexing (LSI)^85^. LSI components 2 through 30 were used for UMAP visualization.

#### Differential expression and accessibility

Differential expression and accessibility analysis was done using the “FindMarkers” function with the “min.pct” parameter set to 0.25 to include only genes or peaks expressed in 25% of cells in either group, comparing between conditions for each cluster independently. Genes and peaks were considered differentially expressed if the Benjamini-Hochberg adjusted p-value was < 0.05 and the log2FC was > ± 0.5.

#### Gene ontology

Significantly upregulated or downregulated genes were analyzed for enrichment by comparison to the GO Biological Process 2025 database in Enrichr^86^. Gene ontology terms were considered significant if the Benjamini-Hochberg adjusted p-value was < 0.05.

#### Motif enrichment analysis

Overrepresented motifs within differentially accessible peaks (DAPs) were identified using the FindMotifs function as part of Signac. DAPs that were not near gene structures were listed as distal intergenic regions. The background set of peaks used for the overrepresentation analysis was derived by matching GC content of the larger set of total open peaks of both karyotypes and the smaller DAP set using the MatchRegionStats function.

#### Histone post translational modification enrichment analysis

Genes lists were analyzed for histone post translational modification (PTM) enrichment by comparison to the ENCODE Histone Modifications 2015 database in Enrichr^86^. For each PTM, the most significant comparison was selected and the PTM was considered significantly enriched if the Benjamini-Hochberg adjusted p-value was < 0.05.

#### Pseudotime

To trace differentiation of NPCs into neurons and astrocyte precursors, pseudotime was performed using Slingshot (version 2.16.0)^87^ with the Cycling NPCs cluster set as the starting cluster. To assess differences in mean pseudotime value between karyotypes, Kolmogorov-Smirnov tests were performed for each lineage. To assess changes in cell density distribution along pseudotime, a general linear regression was performed for each lineage.

#### FigR

FigR (version 0.1.0)^35^ uses the matched single nuclei ATAC-seq and RNA-seq data to define Domains of Regulatory Chromatin (DORCs) driving developmental programs. FigR was performed separately on the Eupl^WC^ and T21^WC^datasets to identify the gene regulatory network driving each karyotype’s developmental program. A cut-off of seven significant peak-gene links was used to define a DORC and a cut-off score of one was used to determine significant transcription factor motif enrichments. To examine changes in DORC accessibility and expression over developmental time, the median accessibility and mean expression per cell for each DORC was calculated and plotted versus the pseudotime value derived from the neural lineages identified via Slingshot. Cells were grouped into 100 pseudotime bins and the rows scaled for clarity. We performed cross correlations between the median accessibility score and mean expression value for each bin to find the maximum correlation value for each DORC in each karyotype.

### CUT&Tag

P3 NPCs derived from the WC isogenic cell line (n=3 biological replicates per karyotype, 500,000 cells per replicate) were collected and cryopreserved. Cryopreserved cells were sent to Active Motif for their Cleavage Under Targets and Tagmentation (CUT&Tag) service, as described previously^88^. Chromatin extraction, tagmentation, precipitation with rabbit anti-H3K27me3 (Active Motif, AB_2561020), and library preparation was performed by Active Motif. Libraries were sequenced on the Illumina NextSeq 550, resulting in 38bp paired-end reads. The Burrows Wheeler Aligner with default settings (mem mode) was used to align reads to the human genome hg38. All duplicate reads were removed, leaving only reads that mapped uniquely (mapping quality>1). Peaks were called using the Model based Analysis of ChIP-seq (MACS) 2.1.0 algorithm at a cut-off of *p*=1e-7. Peaks were used as the input into Active Motif’s proprietary analysis program, which created datasheets containing key data on sample comparisons, peak locations, peak metrics, and gene annotations. For differential analysis between conditions (euploid vs. T21), reads were counted in all merged peak regions and each condition’s replicates were compared using DESeq2. The fold change between euploid and T21 samples was then calculated. Differential peaks with an absolute log_2_ fold change value greater than 1 and p_adj_<0.05 were used for further analysis.

## Resource availability

### Materials availability

No new unique reagents were generated in this study.

### Data and code availability

All sequencing datasets generated in this study have been deposited in the Gene Expression Omnibus (GEO). No new code was generated for this study.

## Acknowledgements

We thank members of the Barrett lab for fruitful discussions and insights into this project. We thank the Harvard University Bauer Sequencing Core for assistance with multiomic analyses and the Dana-Farber Cancer Center Institute Molecular Biology Core Facilities for assistance with bulk transcriptional analyses. We thank the Harvard Department of Stem Cell and Regenerative Biology – Harvard Stem Cell Institute Flow Cytometry Core, and the Broad Technology Space – Flow Cytometry core, in particular Andrew Patentreger, for their assistance with flow cytometry analyses. This work was supported by NIH R01HD111876 to L.E.B.

## Author Contributions

J.A.K. and L.E.B. conceived the study and wrote the manuscript; J.A.K., A.N., S.U., and D.S. generated cells; J.A.K. and S.U. performed data analyses; E.Z., R.M. and W.H. performed proteomic experiments and analyses and advised on experimental design; L.E.B. supervised the study and secured funding. All authors discussed results and contributed to the manuscript.

## Declaration of Interests

The authors declare no competing interests.

**Supplemental Figure 1: Characterization of NPCs produced by dual SMADi**

**Supplemental Figure 2: Astrocyte precursors express canonical markers**

**Supplemental Figure 3: tet-O-GFP clonal analyses and assessment of apoptosis and senescence**

**Supplemental Figure 4: NPC proliferation assessed by flow cytometry and immunocytochemistry**

**Supplemental Figure 5: Alterations in the G1 / S ratio in T21^DS^ NPCs**

**Supplemental Figure 6: Tandem Mass Tag Proteomics identifies proteins with differential abundance in T21 NPCs**

**Supplemental Figure 7: snRNA-seq quality control metrics**

**Supplemental Figure 8: snATAC-seq quality control metrics**

**Supplemental Figure 9: Additional snRNA-seq and snATAC-seq NPC cluster analysis**

**Supplemental Figure 10: Dysregulation of HSA21 encoded genes and proteins**

**Supplemental Figure 11: Altered neurogenic signal in T21 proteome**

**Supplemental Figure 12: Kolmogorov-Smirnov tests identify significant differences in mean pseudotime values**

**Figure 13: Separate molecular program in T21^WC^ driven by distinct DORCs and transcriptional network**

**Supplemental Figure 14: Dysregulated fate-instructive genes in T21 are enriched for heterochromatin marks**

**Supplemental Figure 15: H3K27me3 CUT&Tag QC Metrics**

**Supplemental Figure 16: Additional EZH2i treatment conditions**

**Supplemental Table S1: NPC Differential Protein Abundance**

**Supplemental Table S2: snRNA-seq DEGs**

**Supplemental Table S3: snATAC-seq DAPs**

**Supplemental Table S4: Differential H3K27me3 Peaks**

**Supplemental Table S5: RNA-seq DEGs after GSK126**

## Notes

### Competing Interest Statement

The authors have declared no competing interest.

