## Supplemental Figures for "Aberrant chromatin remodeling influences human neural cell fate change in Trisomy 21"

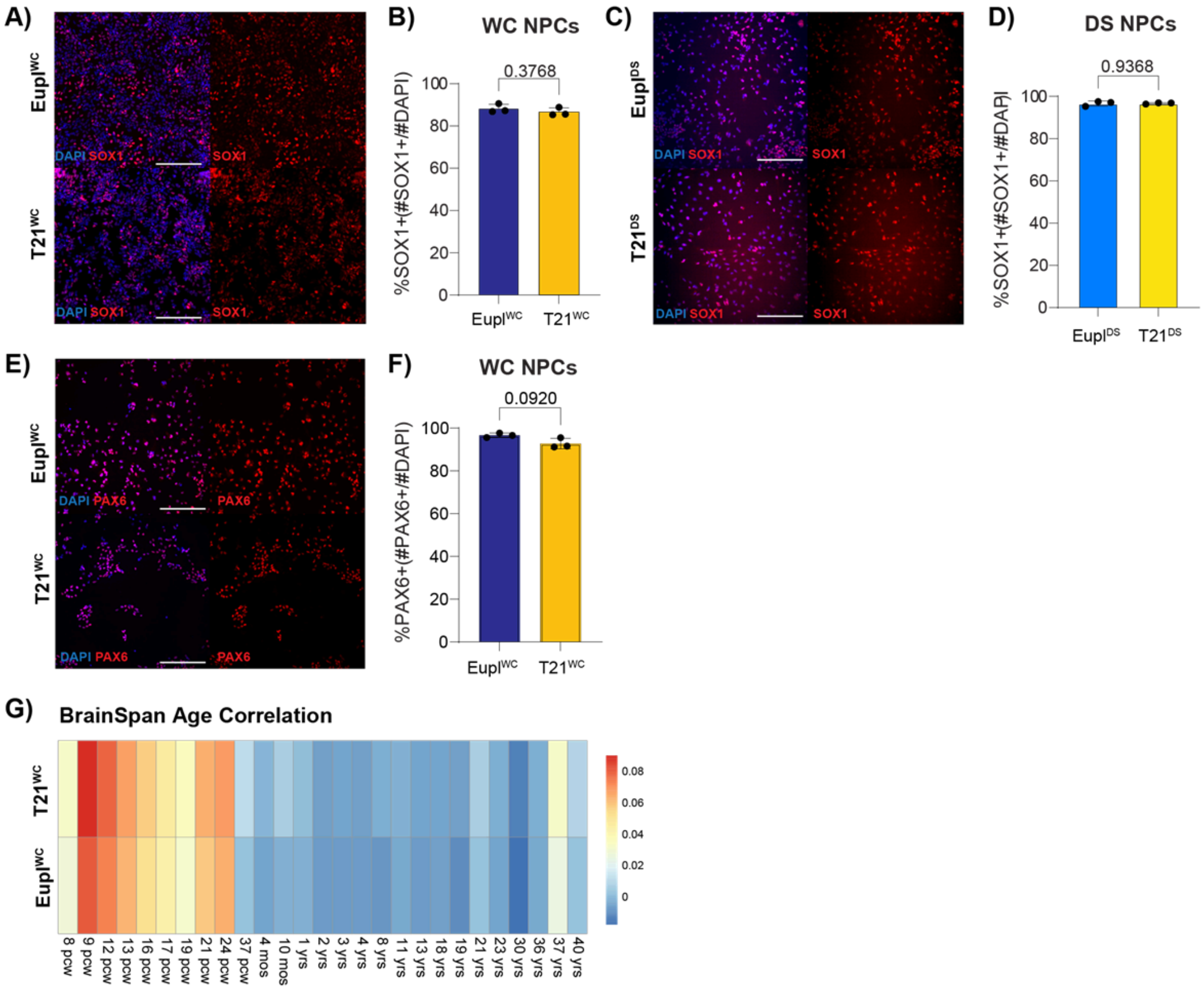


**Supplemental Figure 1: Characterization of NPCs produced by dual SMADi**

**A)** Representative immunocytochemistry of Eupl$^{WC}$/T21$^{WC}$ P3 NPCs immunostained for SOX1 and counterstained with DAPI. **B)** Quantification of SOX1 positive cells shows no difference in the percentage of NPCs produced between Eupl$^{WC}$ and T21$^{WC}$ (p=0.3768). **C)** Representative immunocytochemistry of Eupl$^{DS}$/T21$^{DS}$ P3 NPCs immunostained for SOX1 and counterstained with DAPI. **D)** Quantification of SOX1 positive cells shows no difference in the percentage of NPCs between Eupl$^{DS}$ and T21$^{DS}$ (p=0.9368). **E)** Representative immunocytochemistry of Eupl$^{WC}$/T21$^{WC}$ P3 NPCs immunostained for PAX6 and counterstained with DAPI. **F)** Quantification of PAX6 positive cells shows no difference in the percentage of NPCs produced between Eupl$^{WC}$ and T21$^{WC}$ (p=0.0920). Significance was determined by Welch's t test. Scale bar = 200nm. **G)** Comparison to the BrainSpan transcriptional atlas shows that the NPCs most closely match human prenatal time-points. Scale bar = Pearson correlation value.

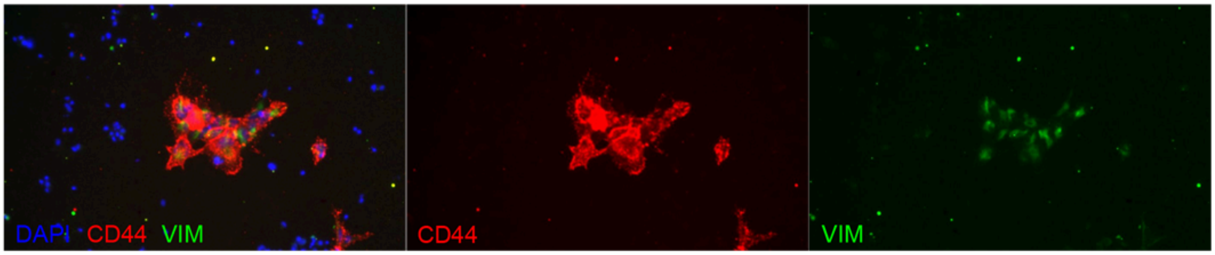


**Supplemental Figure 2 - Astrocyte precursors express canonical marks**
Following spontaneous differentiation of NPCs, astrocyte precursor cells co-express canonical markers CD44 and Vimentin (VIM).

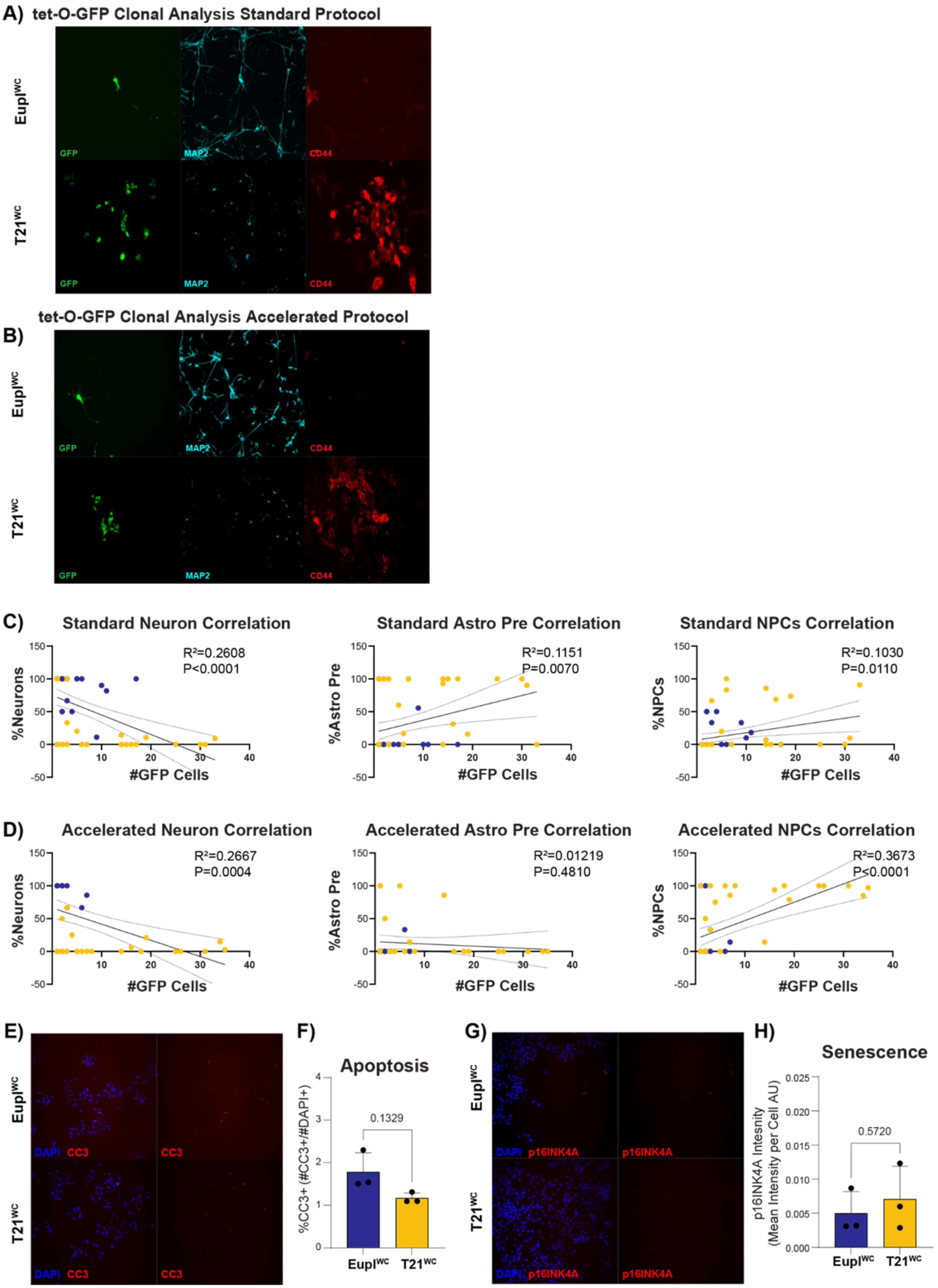
A) tet-O-GFP Clonal Analysis Standard Protocol
EuplWC
T21WC
GFP
MAP2
CD44
tet-O-GFP Clonal Analysis Accelerated Protocol
B)
C) Standard Neuron Correlation
R²=0.2608
P<0.0001
%Neurons
#GFP Cells
Standard Astro Pre Correlation
R²=0.1151
P=0.0070
%Astro Pre
Standard NPCs Correlation
R²=0.1030
P=0.0110
%NPCs
D) Accelerated Neuron Correlation
R²=0.2667
P=0.0004
Accelerated Astro Pre Correlation
R²=0.01219
P=0.4810
Accelerated NPCs Correlation
R²=0.3673
P<0.0001
E)
DAPI CC3
CC3
F) Apoptosis
%CC3+ (#CC3+/#DAPI+)
0.1329
G)
DAPI p16INK4A
p16INK4A
H) Senescence
p16INK4A Intesity (Mean Intensity per Cell AU)
0.5720

**Supplemental Figure 3: tet-O-GFP clonal analyses and assessment of apoptosis and senescence**

**A-B)** Individual images from dual infection of $Eupl^{WC}$ and $T21^{WC}$ iPSCs with rtTA and tet-O-GFP to identify the progeny (GFP+) of single NPCs including neurons (MAP2+) and astrocyte precursors (CD44+) after standard (A) or 1 week accelerated (B) spontaneous differentiation. Merged image shown in Main Fig. 1. **C)** Correlation analyses in the standard time course showing negative correlation between the number of GFP+ cells and the percentage of neurons per clone ($R^2$=0.2608, p<0.0001), a positive correlation between the number of GFP+ cells and the percentage of astrocyte precursors per clone ($R^2$=0.1151, p=0.0070), and a positive correlation between the number of GFP+ cells and the percentage of NPCs per clone ($R^2$=0.1030, p<0.0001). **D)** Correlation in the accelerated time course showing a negative correlation between the number of GFP+ cells and percentage of neurons per clone ($R^2$=0.2667, p<0.0001), no significant correlation between the number of GFP+ cells and the percentage of astrocyte precursors per clone ($R^2$=0.01219, p=0.4810), and a positive correlation between the number of GFP+ cells and the percentage of NPCs per clone ($R^2$=0.3673, p<0.0001). **E)** Representative immunocytochemistry of CC3, a marker of apoptosis, in P3 $Eupl^{WC}$ and $T21^{WC}$ NPCs. **F)** Quantification showing no significant difference in the percentage of CC3+ cells between karyotypes (p=0.1329). **G)** Representative immunocytochemistry of p16INK4A, a marker of senescence, in P3 $Eupl^{WC}$ and $T21^{WC}$ NPCs. **H)** Quantification showing no significant difference in the mean intensity of p16INK4A immunostaining between karyotypes at this time-point (p=0.5720). Significance for F) and H) was determined by Welch's t test.

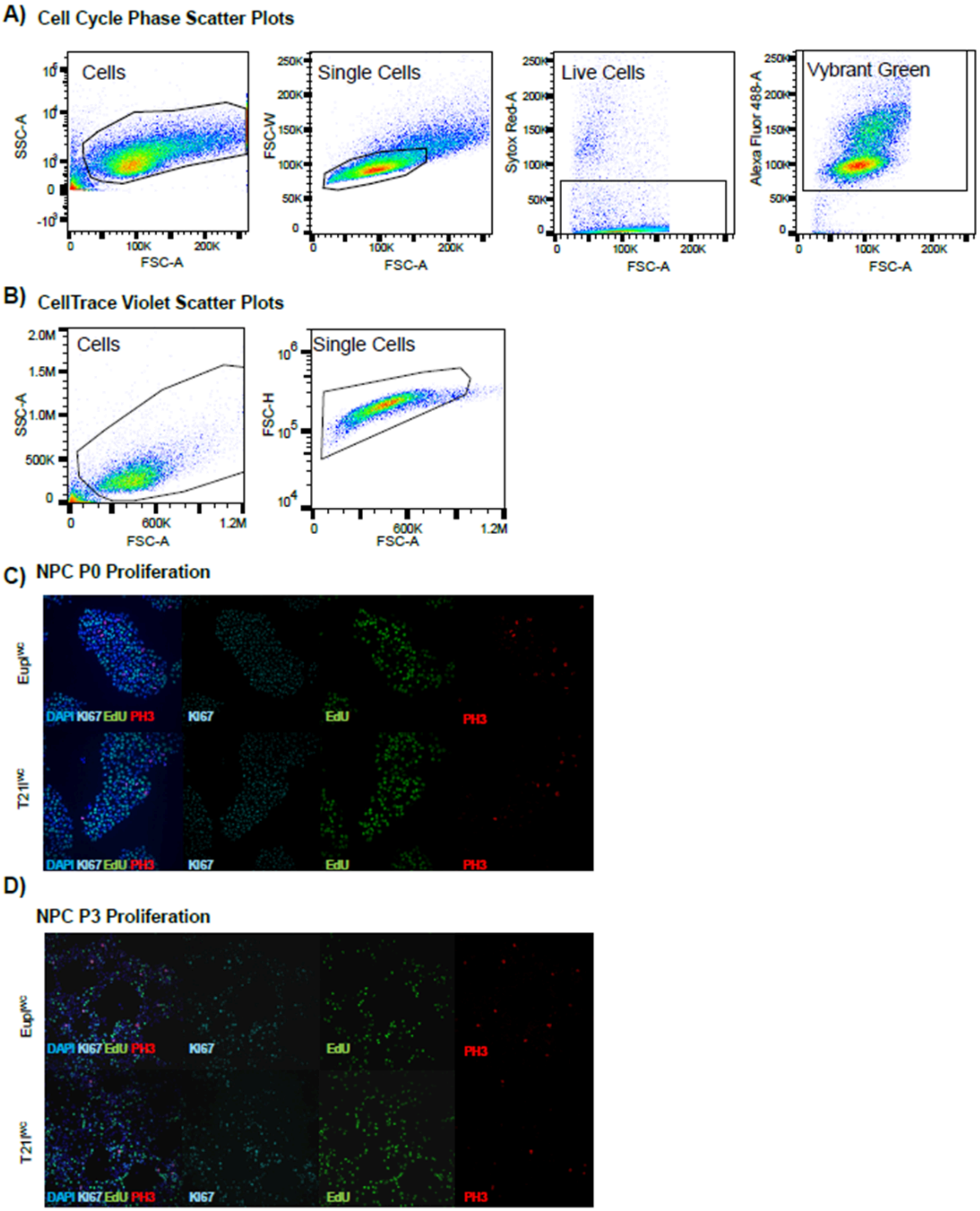


**Supplemental Figure 4: NPC proliferation assessed by flow cytometry and immunocytochemistry**
**A)** Representative flow cytometry scatter plots of cells labeled with Sytox Red and Vybrant DyeCycle Green. **B)** Representative flow cytometry scatter plots of cells labeled with CellTrace Violet. **C)** Representative immunocytochemistry of $Eupl^{WC}$ and $T21^{WC}$ P0 NPCs to identify all proliferating cells (Ki67), cells in S phase (EdU), and cells in G2M (PH3). **D)** Representative immunocytochemistry of $Eupl^{WC}$ and $T21^{WC}$ P3 NPCs to identify all proliferating cells (Ki67), cells in S phase (EdU), and cells in G2M (PH3).

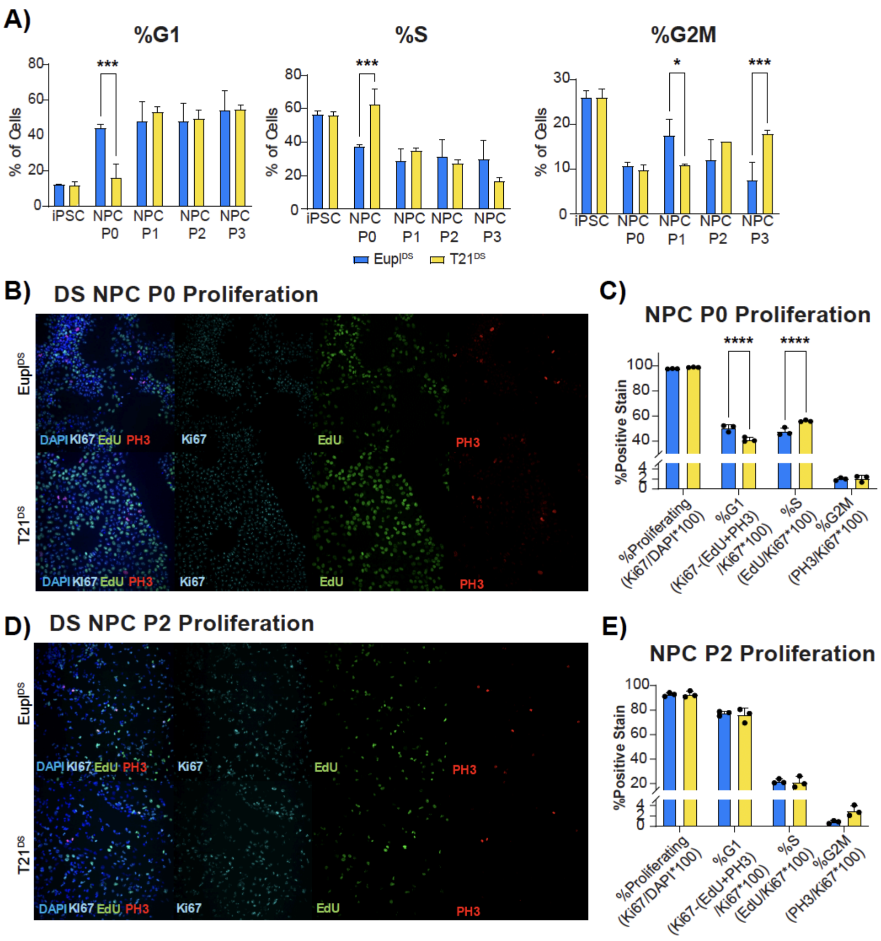


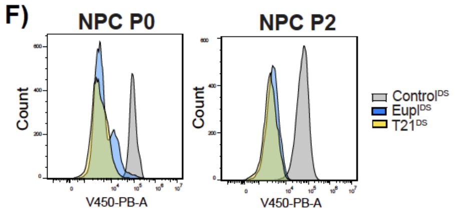


G)

| | Relative Proliferation | P-Value |
|---|---|---|
| DS P0 | 2.11 | 0.00203 |
| DS P2 | 1.20 | 0.000432 |

**Supplemental Figure 5: Alterations in the G1 / S ratio in T21$^{DS}$ NPCs**

**A)** Quantification of dye intensity identified significant shifts in all cell cycle phases between karyotypes. At P0, there was a significant decrease in the proportion of NPCs in G1 in T21$^{DS}$ (44.3 ± 2.1% Eupl$^{DS}$ vs 16.5 ± 7.2% in T21$^{DS}$ $p<0.0001$) with a concurrent increase in the proportion of NPCs in S phase (37.3 ± 1.5% Eupl$^{DS}$ vs 62.7 ± 9.0% in T21$^{DS}$ $p<0.0001$). Additionally, significant changes in the proportion of cells in G2M were identified (P1=0.0149), P3=0.0002). **B)** P0 NPCs immunostained for proliferation markers to identify all dividing cells (Ki67), cells in S phase (EdU), and cells in G2/M (PH3). **C)** Quantification showings a significant increase in the proportion of cells in S phase in T21$^{DS}$ P0 NPCs compared to Eupl$^{DS}$ (47.5 ± 3.1% Eupl$^{DS}$ vs 56.1 ± 0.7% in T21$^{DS}$ $p<0.0001$) along with a decrease in the proportion of NPCs in G1 in T21$^{DS}$ (50.4 ± 2.7% Eupl$^{DS}$ vs 41.2 ± 1.8% in T21$^{DS}$ $p<0.0001$). **D)** At P2, NPCs were again immunostained for proliferation markers that identified all dividing cells (Ki67), cells in S phase (EdU), and cells in G2/M (PH3) **E)** Quantification at P2 showing no significant differences in the percentage of cells proliferating ($p>0.9999$) or in the proportion of cells in different phases of the cell cycle (G1: p=0.9557; S: p=0.9969; G2M: p=0.8627). **F)** Representative traces of flow cytometry analysis of CellTrace Violet showing that **G)** at P0 T21$^{DS}$ NPCs proliferated at a rate 2.11x faster than Eupl$^{DS}$ (p=0.00203) and at P2 T21$^{DS}$ NPCs proliferated at a rate 1.20x faster than Eupl$^{DS}$ ($p=4.32\times10^{-4}$). For A), C), and E) significance was determined by Two-way ANOVA followed by Sidak test for multiple comparisons. For G), significance was determined by Welch's t test.

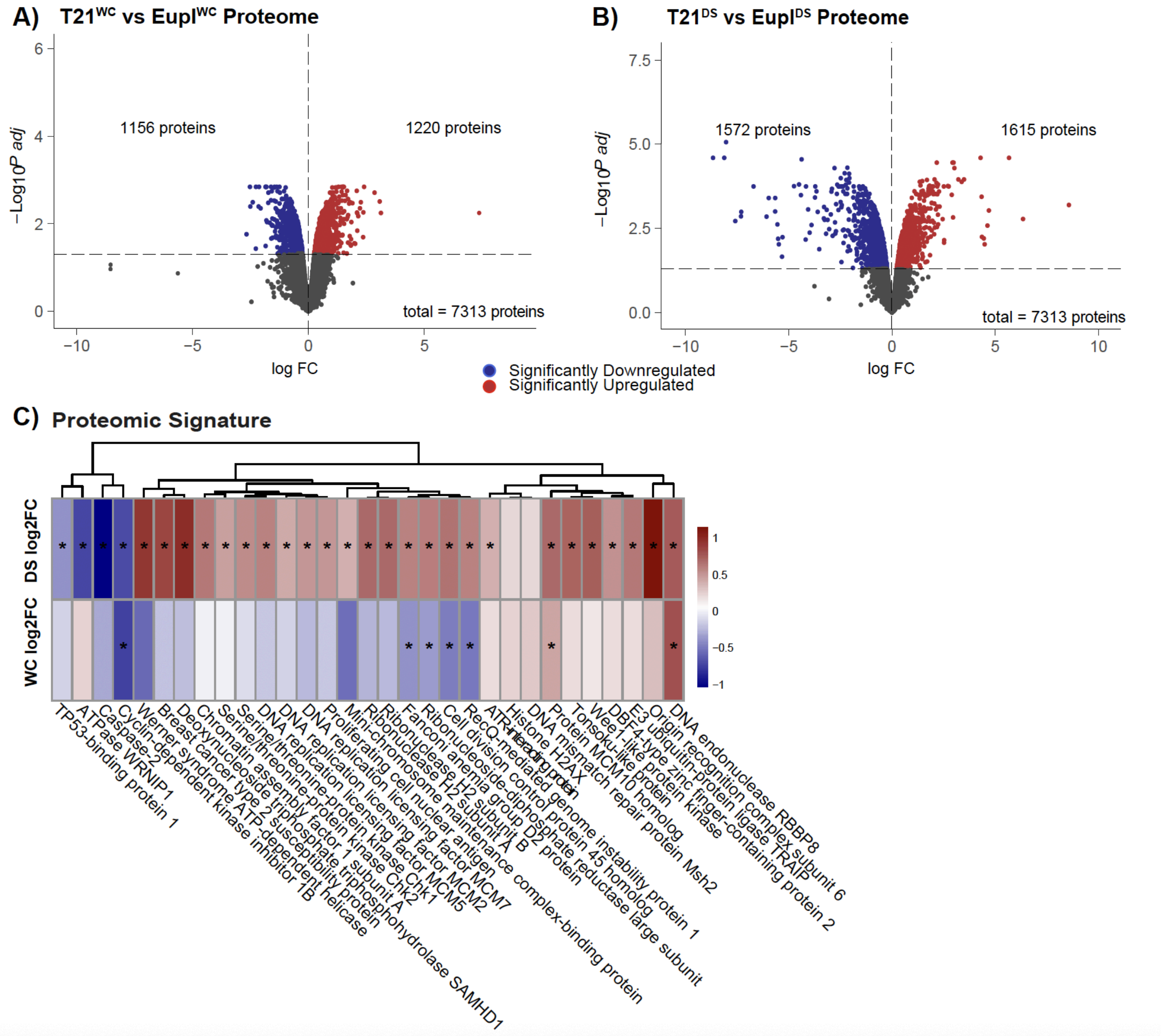


**Supplemental Figure 6: Tandem Mass Tag Proteomics identifies proteins with differential abundance in T21 NPCs**

Mass spectrometry based whole-proteome analyses of P3 NPCs detected 7313 proteins in each sample. **A)** There were 1156 significantly downregulated proteins and 1220 significantly upregulated proteins in T21$^{WC}$ compared to Eupl$^{WC}$ and **B)** 1572 significantly downregulated proteins compared to 1615 upregulated proteins in T21$^{DS}$ compared to Eupl$^{DS}$. **C)** T21$^{DS}$ had multiple significantly upregulated proteins associated with proliferation compared with Eupl$^{DS}$, and in contrast to T21$^{WC}$.

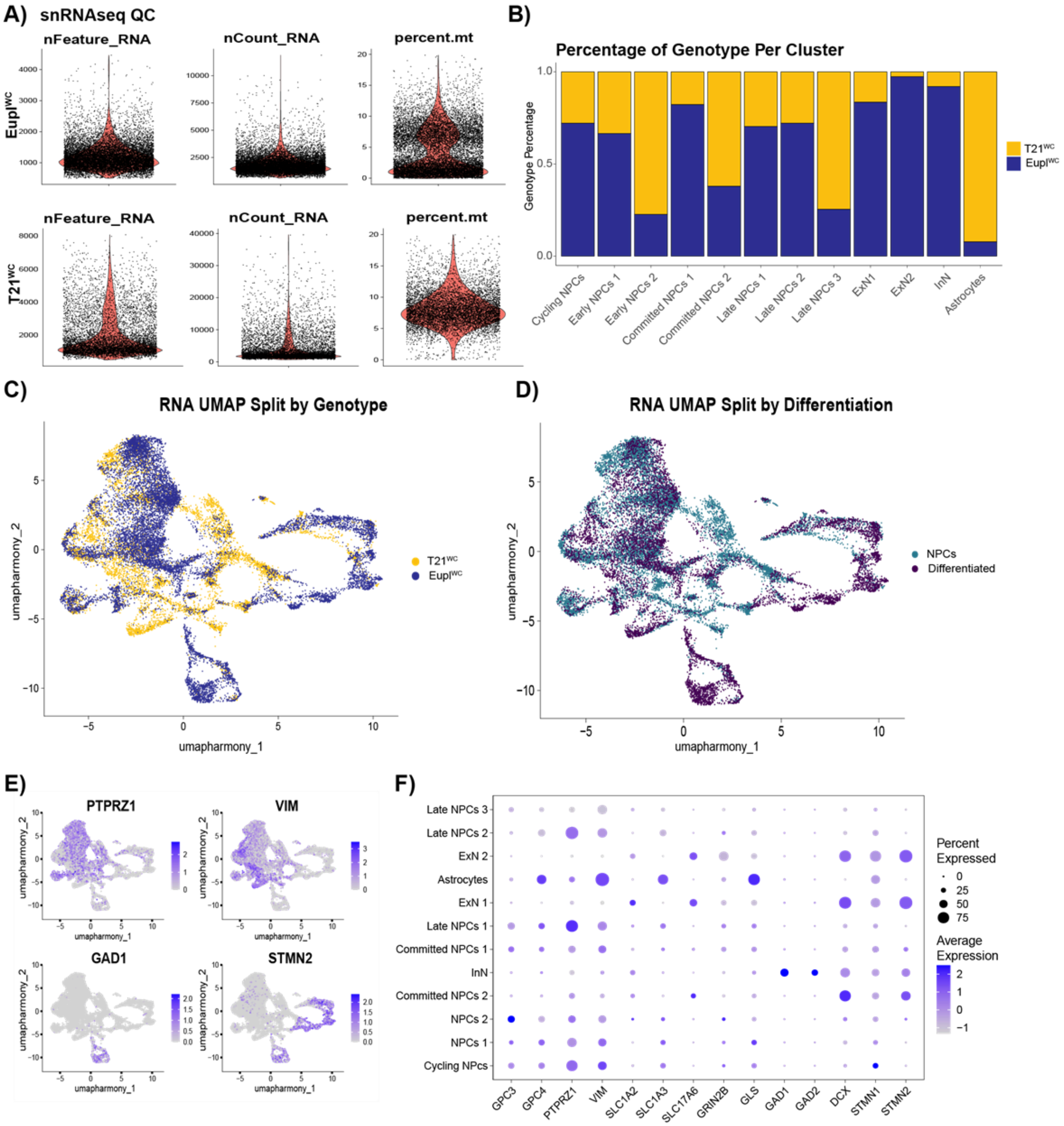


**Supplemental Figure 7: snRNA-seq quality control metrics**

**A)** Quality control metrics of the T21$^{WC}$ and Eupl$^{WC}$ snRNA-seq including the number of detected genes per cell (nFeature_RNA), number of unique molecular identifiers (UMIs) per cell (nCount_RNA), and percent mitochondrial reads (percent.mt). **B)** Percentage of cells per genotype in each cluster. **C)** snRNA-seq UMAP split by karyotype. **D)** snRNA-seq UMAP split by differentiation status. **E)** Selected marker genes used to determine cluster cellular identity overlaid on the snRNA-seq UMAP. **F)** Dotplot showing marker genes used to determine cluster cell type identity.

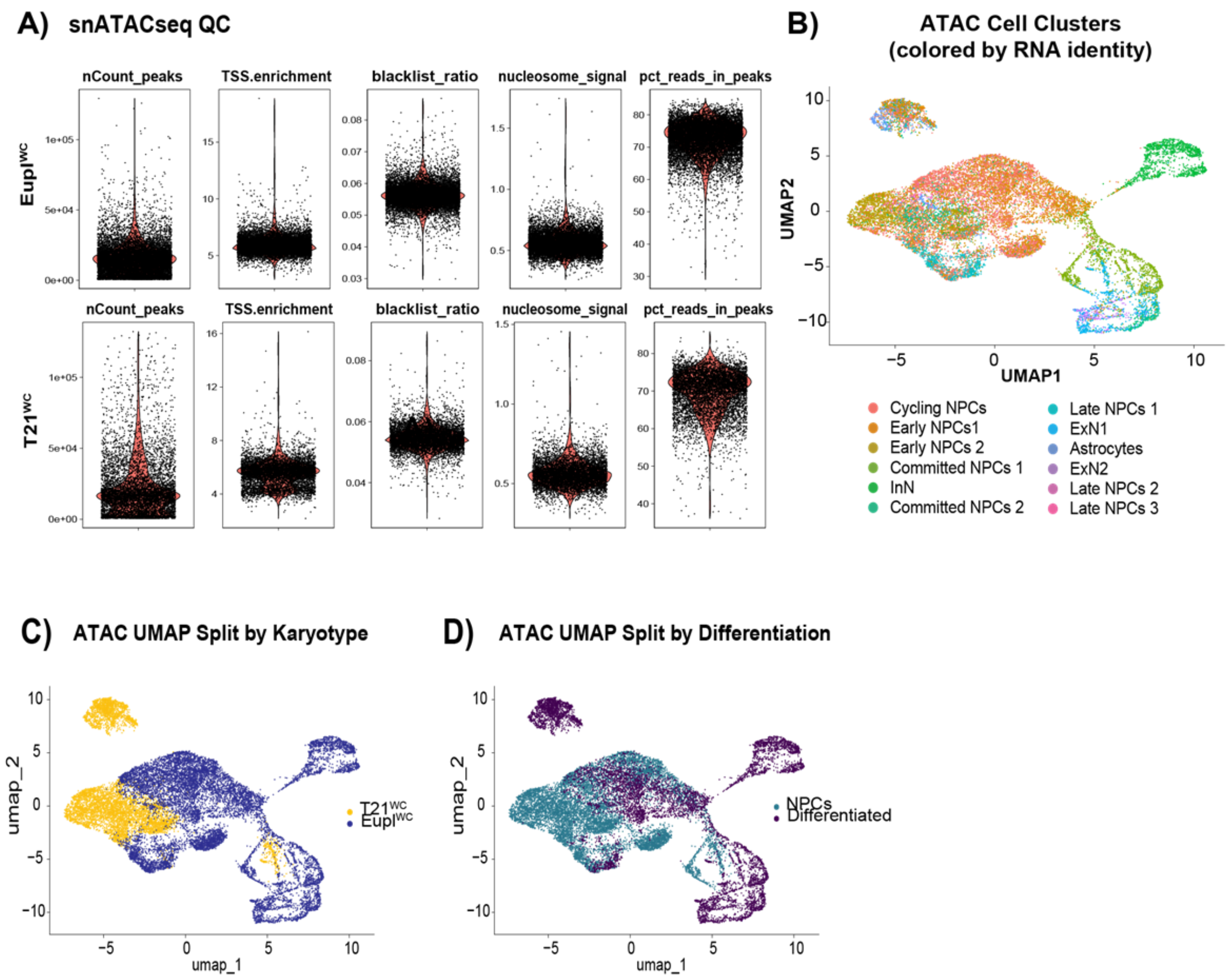


**Supplemental Figure 8: snATAC-seq quality control metrics**

**A)** Quality control metrics of the T21$^{WC}$ and Eupl$^{WC}$ snATAC-seq including number of total peaks (nCount_peaks), enrichment within transcription start sites (TSS.enrichment), ratio of reads in genomic blacklist regions (blacklist_ratio), nucleosome banding pattern (nucleosome_signal), and fraction of reads in peaks (pct_reads_in_peaks). **B)** ATAC-seq UMAP colored by cell clusters identified by transcriptional profile. **C)** ATAC-seq UMAP split by karyotype and **D)** ATAC-seq UMAP split by differentiation status.

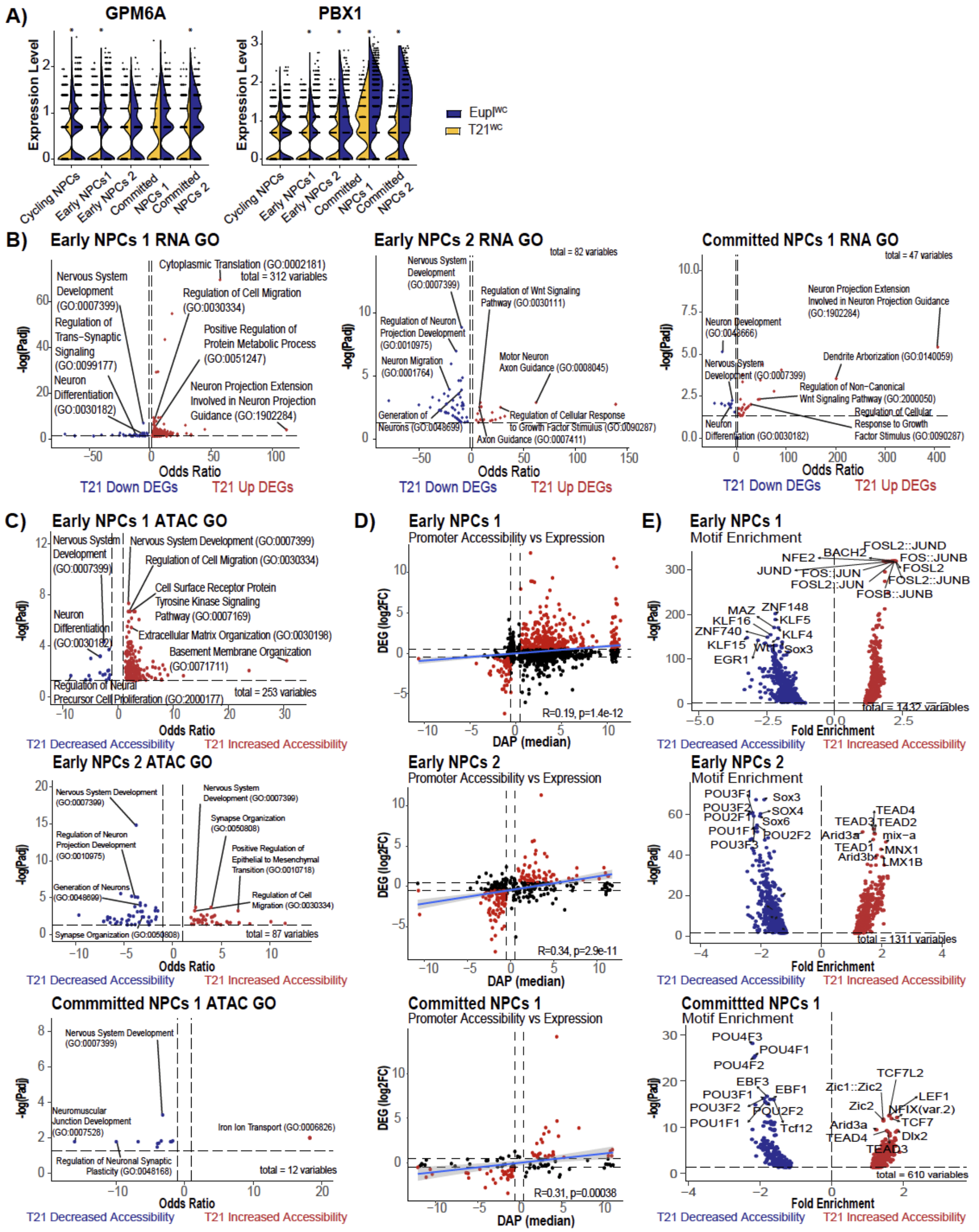


**Supplemental Figure 9: Additional snRNA-seq and snATAC-seq NPC cluster analysis**

**A)** Differential expression analysis identified multiple DEGs known to play a role in neuronal fate such as *GPM6A* and *PBX1*. Significance was determined by the two-sided Wilcoxon Rank-Sums test followed by Benjamini and Hochberg correction for multiple comparisons. **B)** Gene ontology analysis identified enrichment of neurogenesis related terms in the downregulated T21$^{WC}$ DEGs across NPC clusters. **C)** Gene ontology analysis identified enrichment of neurogenesis related terms in the T21$^{WC}$ DAPs with decreased promoter accessibility across NPC clusters. **D)** There were modest but significant correlations determined by general linear regression between differential expression and differential promoter accessibility in each of the NPC clusters (Early NPCs 1: R=0.19, p=1.4e-12; Early NPCs 2: R=0.34, p=2.9e-11; Committed NPCs 1: R=0.31, p=0.00038). **E)** Motif enrichment analysis of distal intergenic regions showed restructuring of the regulatory landscape in T21$^{WC}$ NPCs. Regions with increased accessibility were enriched for pro-proliferative factors and regions with decreased accessibility were enriched with motifs for transcription factors associated with fate specification and neural development.

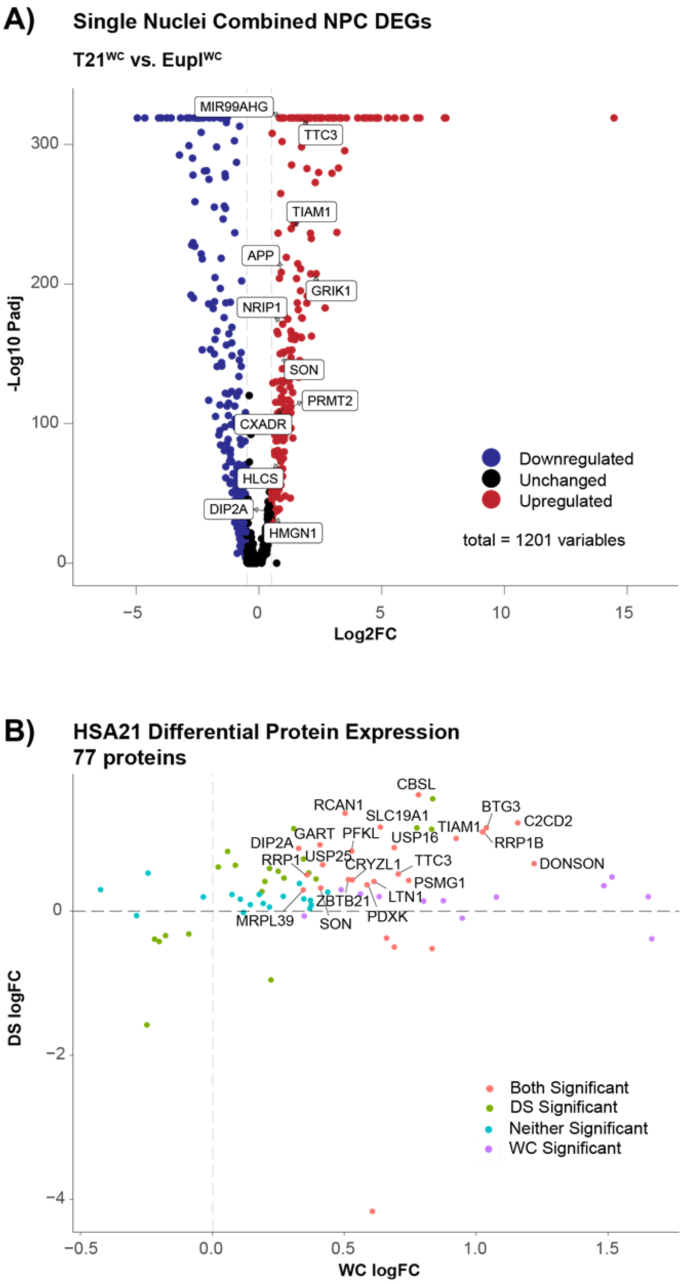


**Supplemental Figure 10: Dysregulation of HSA21-encoded genes and proteins**

**A)** There were 242 unique genes significantly downregulated and 233 unique genes significantly upregulated in combined T21$^{WC}$ NPC clusters compared to Eupl$^{WC}$. Of the upregulated genes, twelve were encoded on HSA21 (labeled above). None of the significantly downregulated genes were encoded on HSA21. **B)** Of 77 proteins from HSA21 detected, 22 were significantly upregulated in both T21$^{WC}$ and T21$^{DS}$ NPCs (labeled above); none were significantly downregulated in both T21$^{WC}$ and T21$^{DS}$ NPCs.

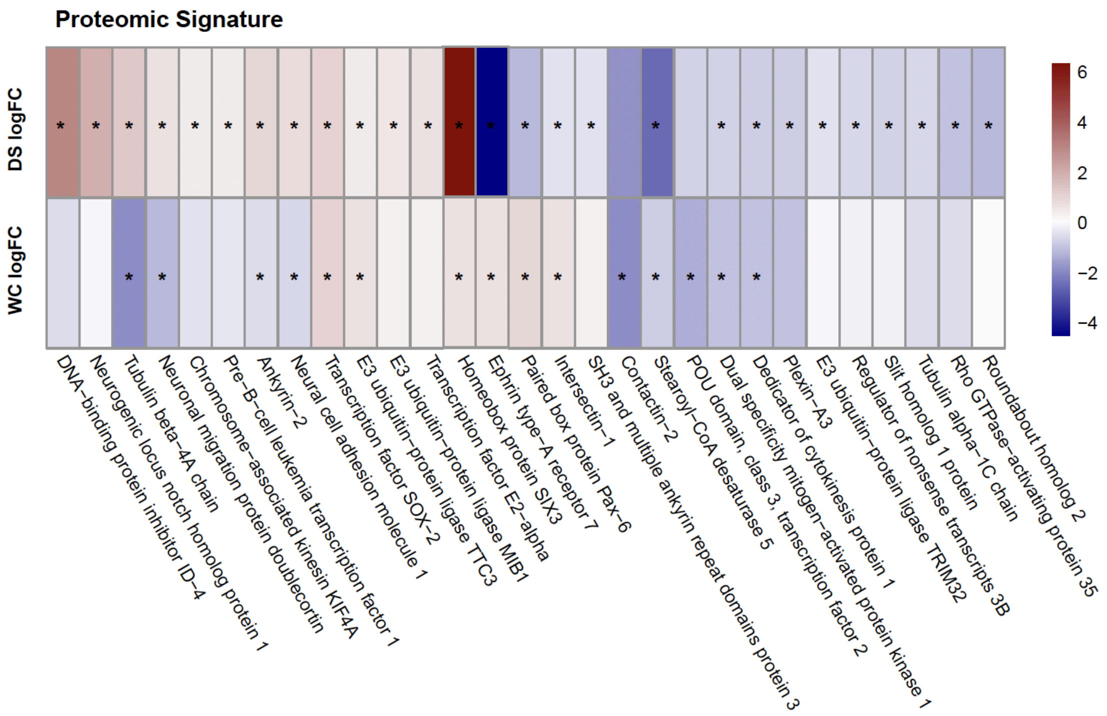


**Supplemental Figure 11: Altered neurogenic signal in T21 proteome**

Whole-proteome analyses of P3 NPCs also revealed dysregulation of neurogenic proteins in both the T21$^{WC}$ and T21$^{DS}$ conditions, relative to their isogenic euploid controls.

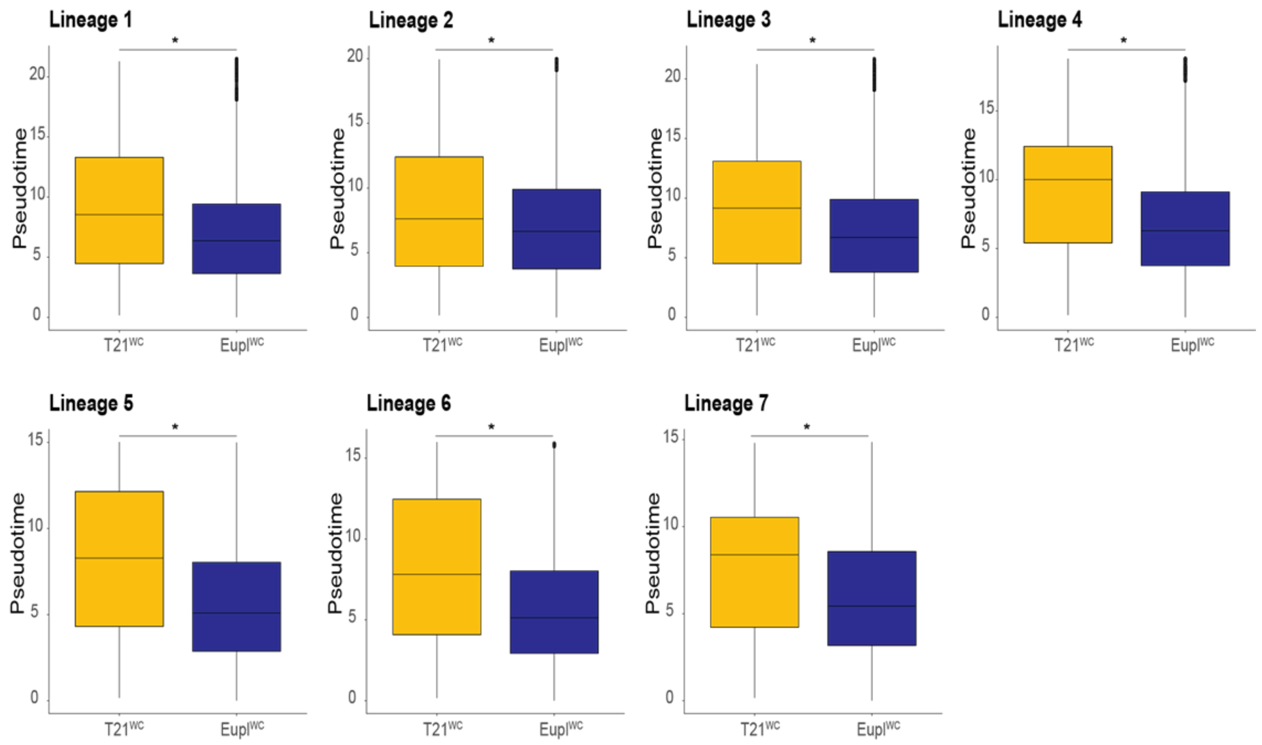


**Supplemental Figure 12: Kolmogorov-Smirnov tests identify significant differences in mean pseudotime values**

Kolmogorov-Smirnov tests resulted in statistically significant differences in each of the seven lineages, with the $T21^{WC}$ condition consistently showing a higher mean pseudotime value compared to $Eupl^{WC}$. P-values for all lineages were $< 2.2x10^{-16}$

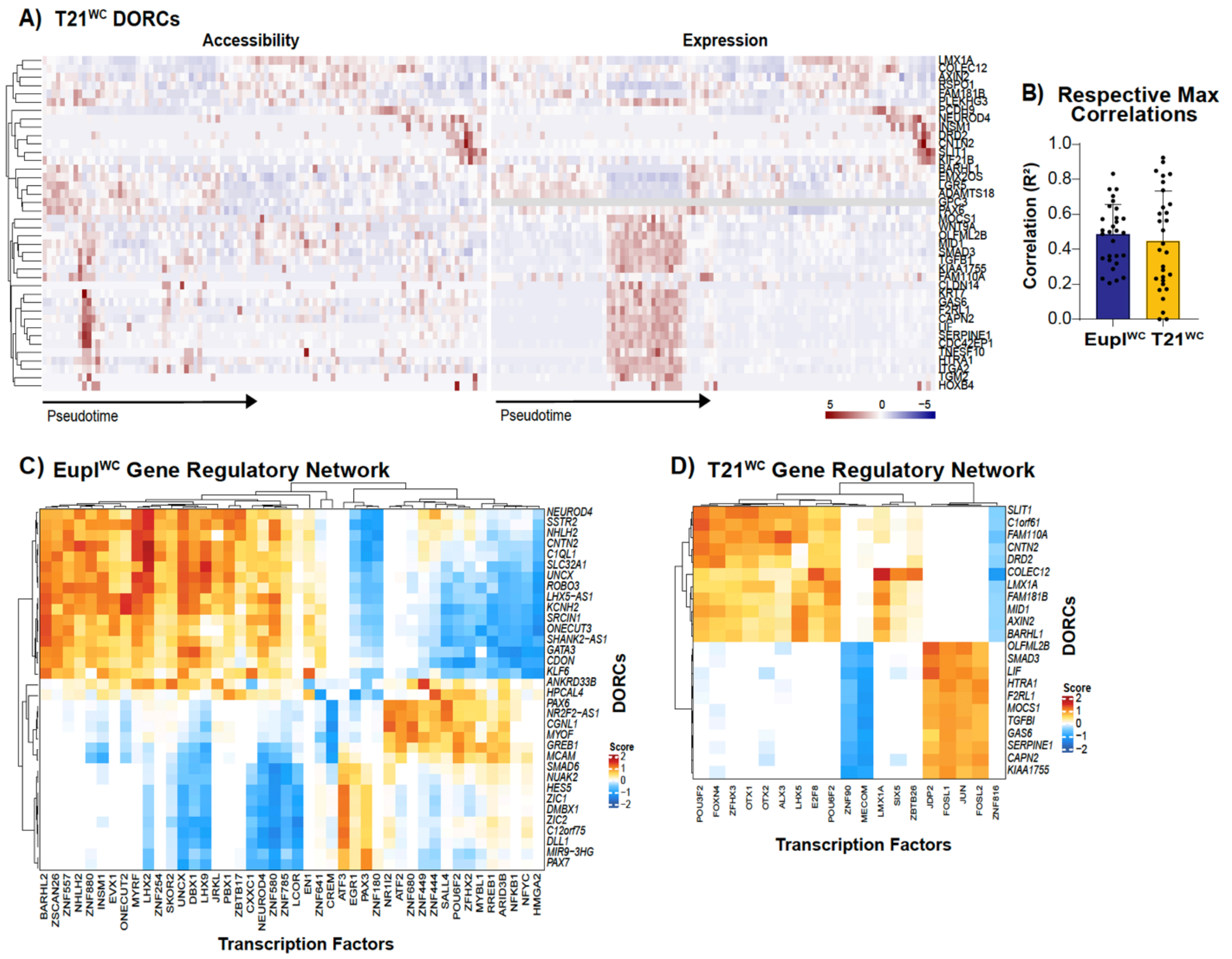


**Supplemental Figure 13: Separate molecular program in T21$^{WC}$ driven by distinct DORCs and transcriptional network**

**A)** A set of 40 DORCs was identified driving the T21$^{WC}$ molecular program with only 3 DORCs overlapping between karyotypes (*CNTN2*, *NEUROD4*, *PAX6*). **B)** There was no significant difference in correlation strength between the respective Eupl$^{WC}$ and T21$^{WC}$ DORCs (p=0.5623). Significance determined by Welch's t test. **C)** Complete network of the 44 transcription factors either activating or repressing the expression of the Eupl$^{WC}$ DORCs. **D)** A separate network of transcription factors controlling DORC expression was identified in T21$^{WC}$.

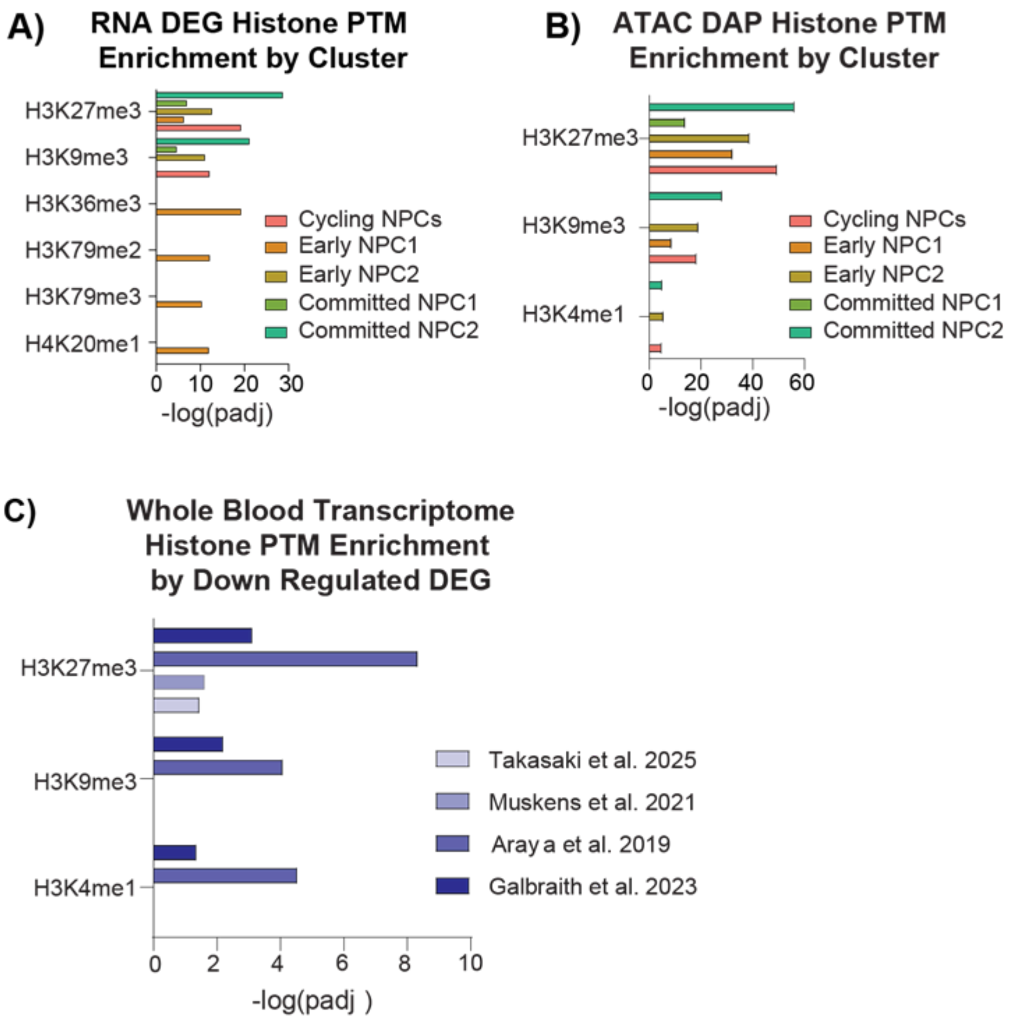


**Supplemental Figure 14: Dysregulated fate-instructive genes in T21 are enriched for heterochromatin marks**
**A)** Comparison of T21WC/EuplWC DEGs to the ENCODE Histone Modifications 2015 database per cluster. **B)** Comparison of T21WC/EuplWC DAPs to the ENCODE Histone Modifications 2015 database per cluster. **C)** In four published blood cell datasets (Takasaki et al. 2025, Muskens et al. 2021, Araya et al. 2019, Galbraith et al. 2023) histone enrichment was only present in the downregulated genes, with H3K27me3 being the top enriched mark across all datasets.

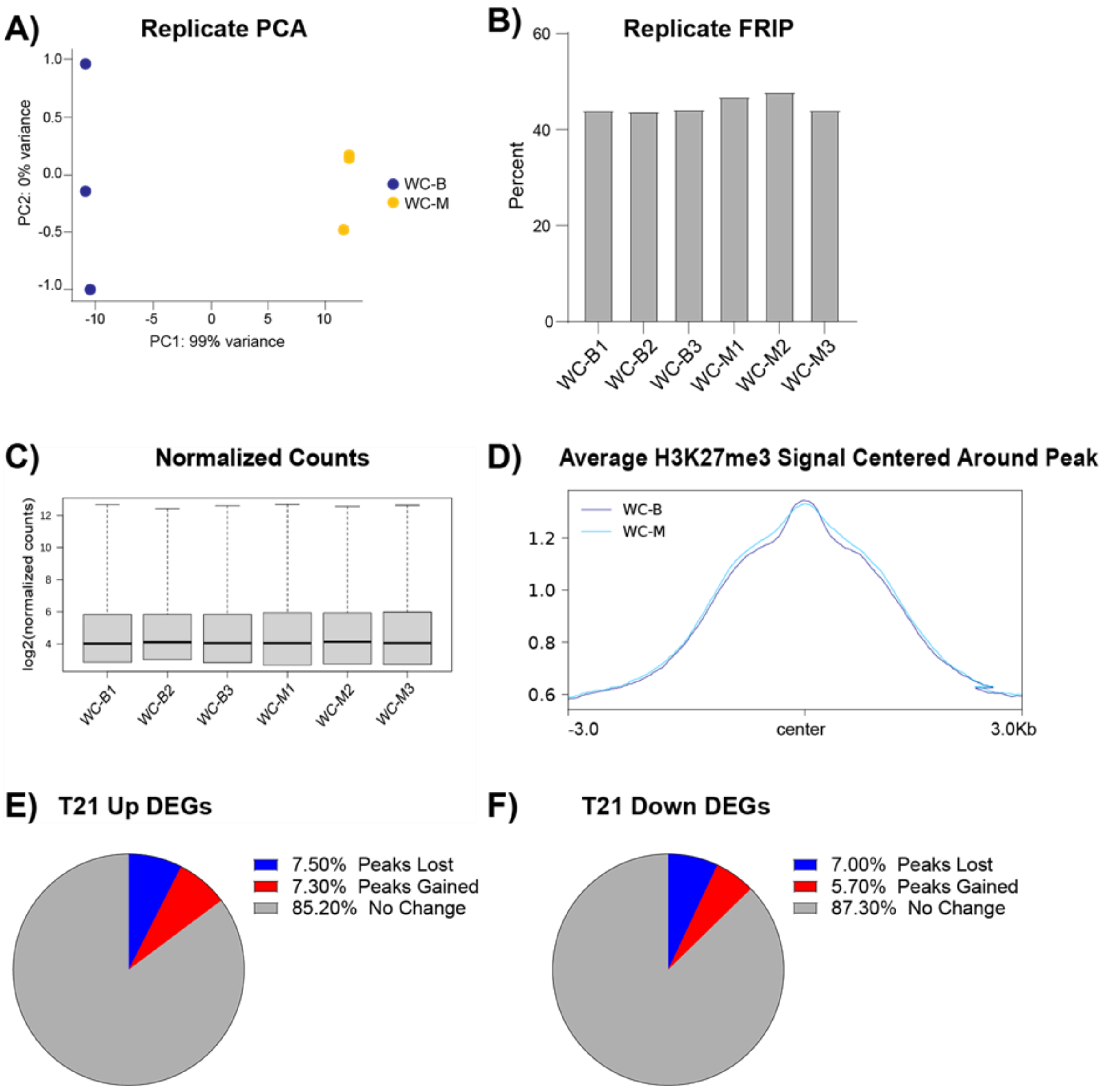


**Supplemental Figure 15: H3K27me3 CUT&Tag QC Metrics**
**A)** PCA plot for Eupl$^{WC}$ and T21$^{WC}$ P3 NPC replicates. **B)** Fraction Read in Peak (FRIP) for each replicate. **C)** Normalized read count for each replicate. **D)** Average H3K27me3 signal around the identified peaks from the Eupl$^{WC}$ and T21$^{WC}$ P3 NPCs. **E)** Genes that contain gained and lost H3K27me3 peaks in DEGs that are **E)** upregulated and **F)** downregulated in T21$^{WC}$ NPCs.

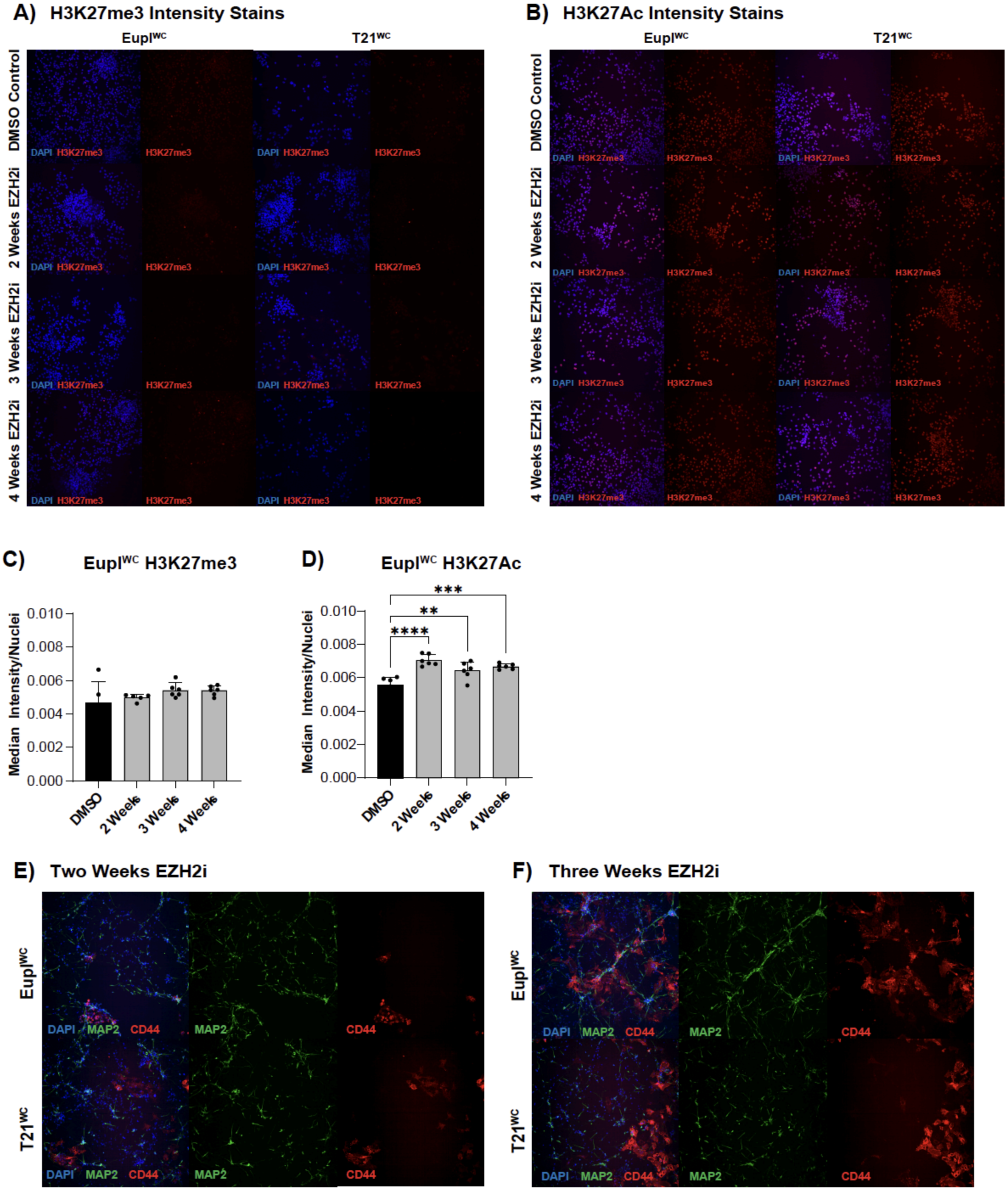


**Supplemental Figure 16: Additional EZH2i treatment conditions**

**A)** Representative immunocytochemistry for H3K27me3 in Eupl$^{WC}$ and T21$^{WC}$ P3 NPCs. **B)** Representative immunocytochemistry for H3K27ac in Eupl$^{WC}$ and T21$^{WC}$ P3 NPCs. **C)** Eupl$^{WC}$ NPCs treated with EZH2i did not show a significant change in H3K27me3 intensity compared to DMSO controls (2 weeks $p=0.7799$; 3 weeks $p=0.1722$; 4 weeks $p=0.1901$). **D)** Eupl$^{WC}$ NPCs treated with EZH2i showed a significant increase in intensity of H3K27ac (2 weeks $p<0.0001$; 3 weeks $p=0.0032$; 4 weeks $p=0.0003$). Representative immunocytochemistry of the Eupl$^{WC}$ and T21$^{WC}$ cells post spontaneous differentiation after either **E)** 2 weeks or **F)** 3 weeks of EZH2i treatment. Significance was determined by One-way ANOVA followed by Dunnett tests for multiple comparisons.

**Table S1: NPC Differential Protein Abundance**
**Table S2: snRNA-seq DEGs**
**Table S3: snATAC-seq DAPs**
**Table S4: Differential H3K27me3 Peaks**
**Table S5: RNA-seq DEGs after GSK126**
